# In Silico Analysis of Drug Off-Target Effects on Diverse Isoforms of Cervical Cancer for Enhanced Therapeutic Strategies

**DOI:** 10.1101/2023.09.09.556929

**Authors:** Azhar Iqbal, Faisal Ali, Moawaz Aziz, Asad Ullah Shakil, Shanza Choudhary, Adiba Qayyum, Fiza Arshad, Sarah Ashraf, Sheikh Arslan Sehgal, Momina Hussain, Muhammad Sajid

## Abstract

Cervical cancer is a severe medical issue as 500,000 new cases of cervical cancer are identified in the world every year. The selection and analysis of the suitable gene target are the most crucial in the early phases of drug design. The emphasis at one protein while ignoring its several isoforms or splice variations may have unexpected therapeutic or harmful side effects. In this work, we provide a computational analysis of interactions between cervical cancer drugs and their targets that are influenced by alternative splicing. By using open-accessible databases, we targeted 45 FDA-approved cervical cancer drugs targeting various genes having more than two distinct protein-coding isoforms. Binding pocket interactions revealed that many drugs do not have possible targets at the isoform level. In terms of size, shape, electrostatic characteristics, and structural analysis have shown that various isoforms of the same gene with distinct ligand-binding pocket configurations. Our results emphasized the risks of ignoring possibly significant interactions at the isoform level by concentrating just on the canonical isoform and promoting consideration of the impacts of cervical cancer drugs on- and off-target at the isoform level to further research.

## 1. Introduction

In developing countries, cervical cancer is the main reason for cancer-related deaths and years of life loss (1). Several years earlier than the median age at which breast, lung, and ovarian cancers are diagnosed, cervical cancer is commonly diagnosed in one’s fifth decade of life (2). Ninety percent of the 270 000 cervical cancer fatalities in 2015 happened in low- and middle-income countries (LMIC), where mortality is 18 times higher than in developed nations (3). Nearly all cervical cancers are caused by high-risk subtypes of the human papillomavirus (HPV), whereas screening and vaccination programs are effective disease preventive measures for HPV (4). The two most prevalent histological subtypes (squamous cell carcinoma, and adenocarcinoma) account for 70% and 25% of all cervical malignancies, respectively (5, 6). The major decrease in cervical cancer mortality has been attributed to the development and implementation of screening programs (7). Cervical cancer has a poor prognosis following metastasis or recurrence; the 5-year overall survival (OS) rate is about 17% (8). In order to improve the treatment efficacy of cervical cancer, it is crucial to uncover novel therapeutic targets and survival-associated biomarkers.

Major innovations in large-scale multi-omics research provide a unique perspective for the systems biology analysis of the emergence and spread of cancer. HPV contributes to the development of cervical cancer, which is considered a virus-driven malignancy. Early HPV infection may simply be a result of external causes, like changes in the genome would eventually cause cervical epithelial cells to convert into malignant (for example, gene fusion, non-coding RNAs, copy number variation, DNA methylation, and somatic DNA mutations) (9-13). Transcriptome and epigenetic modifications have been the focus of the bulk of previous prospective studies. However, Alternative splicing (AS) in cancer post-transcriptional isoforms hasn’t been thoroughly studied yet.

In eukaryotes, a remarkable biological process known as alternative splicing, which promotes proteome diversity, allows a single gene to express several protein isomers. In humans, where more than 94% of genes are alternatively spliced, the occurrence and properties of alternative splicing are also highly diverse (14-16). This method enables cancer cells to generate abnormal proteins with altered functional domains that promote carcinogenesis (17-19). In malignancies, these domain changes can lead to complicated remodeling and protein-protein interactions. Some essential oncogenic splicing variations have the ability to control tumor epithelial-to-mesenchymal transition and biological processes of cancer stem cell (20). Gene expression is properly controlled to occur in a context-specific way, even if gene isoforms commonly appear to have different, sometimes even opposing functions.

Aberrant isoforms, or spliced variations that cause disease, have the potential to be effective drug targets in addition to serve as significant biomarkers (21, 22). In this study, we primarily focused on cervical cancer and examined whether or not the drugs are effective against the target gene isoforms. In this work, we examined the effectiveness of FDA-approved drugs against the various isoforms of the cervical cancer-related genes. Using structural analysis and the clinical data on the expression of these genes, we curated the drug interaction data for the various isoforms of different genes implicated in cervical cancer and evaluated their effectiveness against isoforms.

## 2. Methods

### 2.1 Collection of genes and their isoforms

We found the genes associated with cervical cancer using COSMIC database (23) which is an online resource of somatically acquired mutations reported in human cancer. There are more than 30 genes that may contribute to cervical cancer shown in Supplementary File 1. Based on the number of patient samples, the top 5 genes out of 30 were selected, and these genes were then used for further analysis. The Ensemble genome database (24) was used to curate the isoforms and protein sequences for these genes. Using COSMIC Mutation ID, the mutations were identified in genes and matched with the variants of each isoform using the Ensemble database.

### 2.2 Curation of drugs-target interaction data

By using the Drug Gene Interaction Database (DGIdb) (25), we curated the FDA Approved drugs for our genes. Through this database, more than 40 drugs that have received FDA approval were found. These drugs were retrieved from the Drug Bank (26) and cheMBL (27).

### 2.3 Sequence analysis of isoforms

To check the conservation of binding pocket in isoforms of the genes, Binding Pockets of the canonical proteins were predicted through the COACH Server (https://zhanggroup.org/COACH/). We found domains from EMBL-EBI InterPro database (28) and aligned these with the sequences of the canonical protein and their isoforms. Using the Bioconductor programme msa, which offers a selection of alignment techniques and produces alignment plots in LaTeX format, we created numerous alignments of sequences. Using the Cluster Omega method included in the msa package, we created an alignment of the binding site sequence with all of the protein isoforms of the same gene.

### 2.4 Isoforms expression in normal and tumors samples

We looked at the clinical data offered by UCSC Xena (29) for cervical cancer patients which is an online resource for analyzing multi-omics, clinical, and phenotypic data. We used UCSC Xena to compare TCGA tumor samples to GTEx normal samples to evaluate whether protein coding isoforms are up-or down-regulated in cervical cancer. The expression of protein isoforms was examined in patient normal samples using GTEx and tumor samples using TCGA, both of which were drawn from the 307 Cervical Cancer Samples that are available in the UCSC Xena database. We also visualized the exon structure of the isoforms to better understand the pattern of alternative splicing in the various isoforms of the genes.

### 2.5 Structure Prediction of Protein Isoforms and Ligand Docking

To better understand the associations between the proteins with their ligands (drugs), we predicted the 3D structures of protein isoforms using a number of tools for structural level study of the different isoforms of the proteins. Protein isoform structures were predicted through the use of the structure prediction tools trRosetta (30), Robetta (31), Swiss-Model (32), and I-TASSER (33). Further, the ERRAT quality factor and the favored region, allowed region, and disabled region in the Ramachandran plot were used to evaluate the predicted structures. After evaluating, We utilized SiteMap53 (34) to determine the drug targets region in those protein isoforms’ 3D structures. Through the use of Chimera 1.15rc, predicted 3D structures for the isoforms were further prepared for docking analysis. We used Pyrex software to investigate the ligand-protein docking analysis, and we took into account a number of drugs that have already been approved for such proteins so that we can check these drugs’ effectiveness against various protein isoforms that are affected by disease. Poses of the Protein-Ligand Complexes were captured for further analyzing the pocket sizes, shapes, and electrostatic surfaces of docked protein isoforms.

### 2.6 Interaction analysis

The Discovery Studio 2021 Client was used to examine protein-ligand complexes. We examined that how the drug, which has a high binding affinity value with the canonical protein, interacts with the different isoforms. Further, we examined the interactions between hydrophobic and hydrogen sites in different docked protein isoforms

## 3. Results

### 3.1 Drugs Target Genes have multiple Isoforms

More than 30 genes linked to cervical cancer were identified to have missense mutations show in Supplementary File 1. Keeping in view the number of the patient samples, we chose five genes for further analysis. We found FDA-approved drugs interactions to analyze the interactions among drug and its target protein isoforms. We were able to retrieve more than 145 entries belonging to 5 distinct genes of Cervical Cancer.

A partial list from a summary table is shown in Table 1. We identified the bulk of the candidate genes had two or even more transcribed spliced variants and protein isoforms Fig. 1.

**Table 1.**
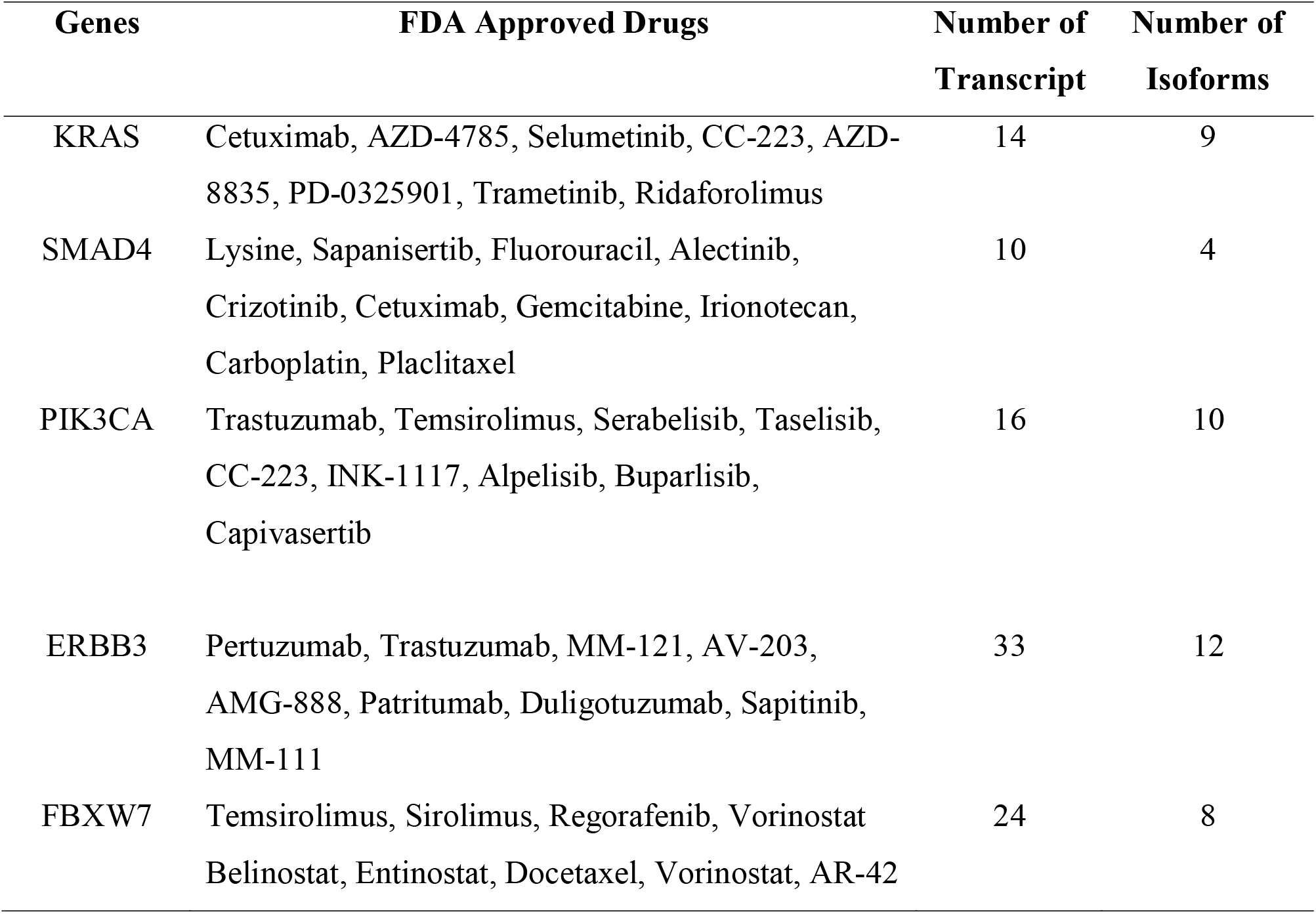
FDA Approved Drugs against target genes and number of protein-coding isoforms.

| Genes | FDA Approved Drugs | Number of Transcript | Number of Isoforms |
| --- | --- | --- | --- |
| KRAS | Cetuximab, AZD-4785, Selumetinib, CC-223, AZD-8835, PD-0325901, Trametinib, Ridaforolimus | 14 | 9 |
| SMAD4 | Lysine, Sapanisertib, Fluorouracil, Alectinib, Crizotinib, Cetuximab, Gemcitabine, Irinotecan, Carboplatin, Paclitaxel | 10 | 4 |
| PIK3CA | Trastuzumab, Temsirolimus, Serabelisib, Taselisib, CC-223, INK-1117, Alpelisib, Buparlisib, Capivasertib | 16 | 10 |
| ERBB3 | Pertuzumab, Trastuzumab, MM-121, AV-203, AMG-888, Patritumab, Duligotuzumab, Sapitinib, MM-111 | 33 | 12 |
| FBXW7 | Temsirolimus, Sirolimus, Regorafenib, Vorinostat, Belinostat, Entinostat, Docetaxel, Vorinostat, AR-42 | 24 | 8 |

**Fig. 1.**
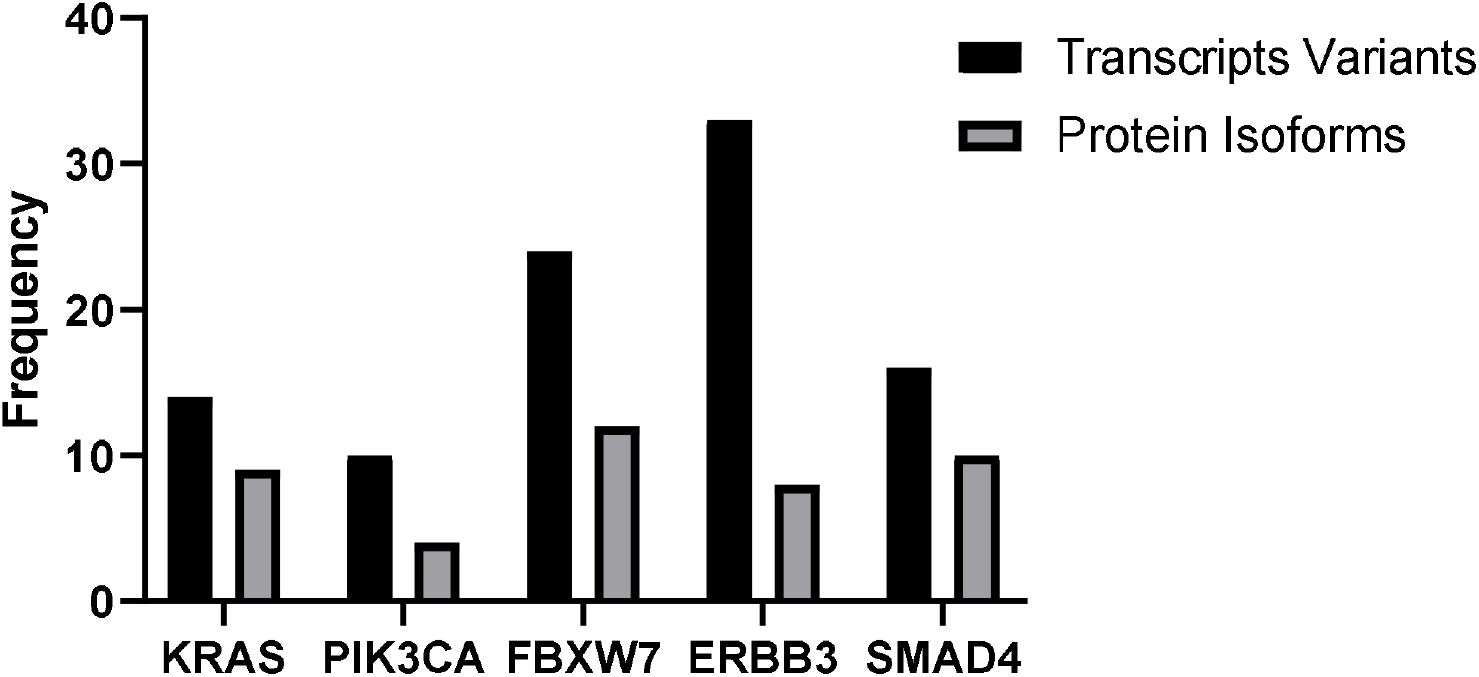
Shows the number of transcript variants and the protein coding isoforms of canonical proteins.

Our findings demonstrate that the majority of cancer drug target genes undergo splicing and make many protein isoforms which may are functionally distinct and react with drugs in different manner, highlighting the significance of getting isoforms and alternative splicing into account in drug development.

### 3.2 Protein isoforms shows differences in the binding pockets

Using several sequence alignments, we were able to pinpoint the precise interaction residues in each isoform’s drug binding region. We carried out multiple sequence alignment between the Pfam functional domains, the canonical protein, isoform sequences, and the predicted protein binding pocket. Here we describe some sequence alignment plots of few genes.

Cellular functions essential for the development of cancer, such as cell growth, proliferation, motility, survival, and metabolism, are regulated by the PI3KCA protein (35). PIK3CA gene has 4 isoforms (PIK3CA-201, PIK3CA-203, PIK3CA-204 and PIK3CA-205). Isoforms PIK3CA-203 & 204 have 21 and 118 residues respectively which completely lacks the predicted pocket binding Fig. 2. Canonical Protein and Isoforms PIK3CA-201 & 205 found to have identical sequences in the predicted binding pocket. However, we found variations in the C-terminal regions and domain PF00454 of isoforms PIK3CA-201 & 205 Fig. 2. We examined the C terminal region of the Canonical protein, PIK3CA-201 & 205, and Pfam domain PF00454 to further explain this variation. Through the previous studies we found that the C terminal region i necessary for catalysis. In the absence of membrane, it reduces the enzyme’s baseline activity while promoting membrane binding. This has been suggested to be a crucial PI3Ks regulating component (36). And the Pfam domain is one of the domains of p100_α_ catalytic subunit of the PIK3CA. However, in USP13-PIK3CA, the whole C-terminal region is replaced with the USP13 protein, which affects catalysis. Since PIK3CA-201 and PIK3CA-205 have the same upstream regions overall, the fusion proteins produced by the two isoforms should ideally have the same structure. Additionally, we aligned two other USP13-PIK3CA protein sequences in the FusionGDB database to support this claim, and all sequences have overlapping interference residues with the predicted pocket binding Supplementary File 2. This sequence-level data indicates that the drug may target all of the USP13-PIK3CA fusion protein’s splice-variant isoforms; as a result, splice-variation within the PIK3CA gene does not influence the binding to its targets in isoforms PIK3CA-201 &205 while it may affect the PIK3CA-203 & 204 which does not have the predicted binding pocket.

**Fig. 2.**
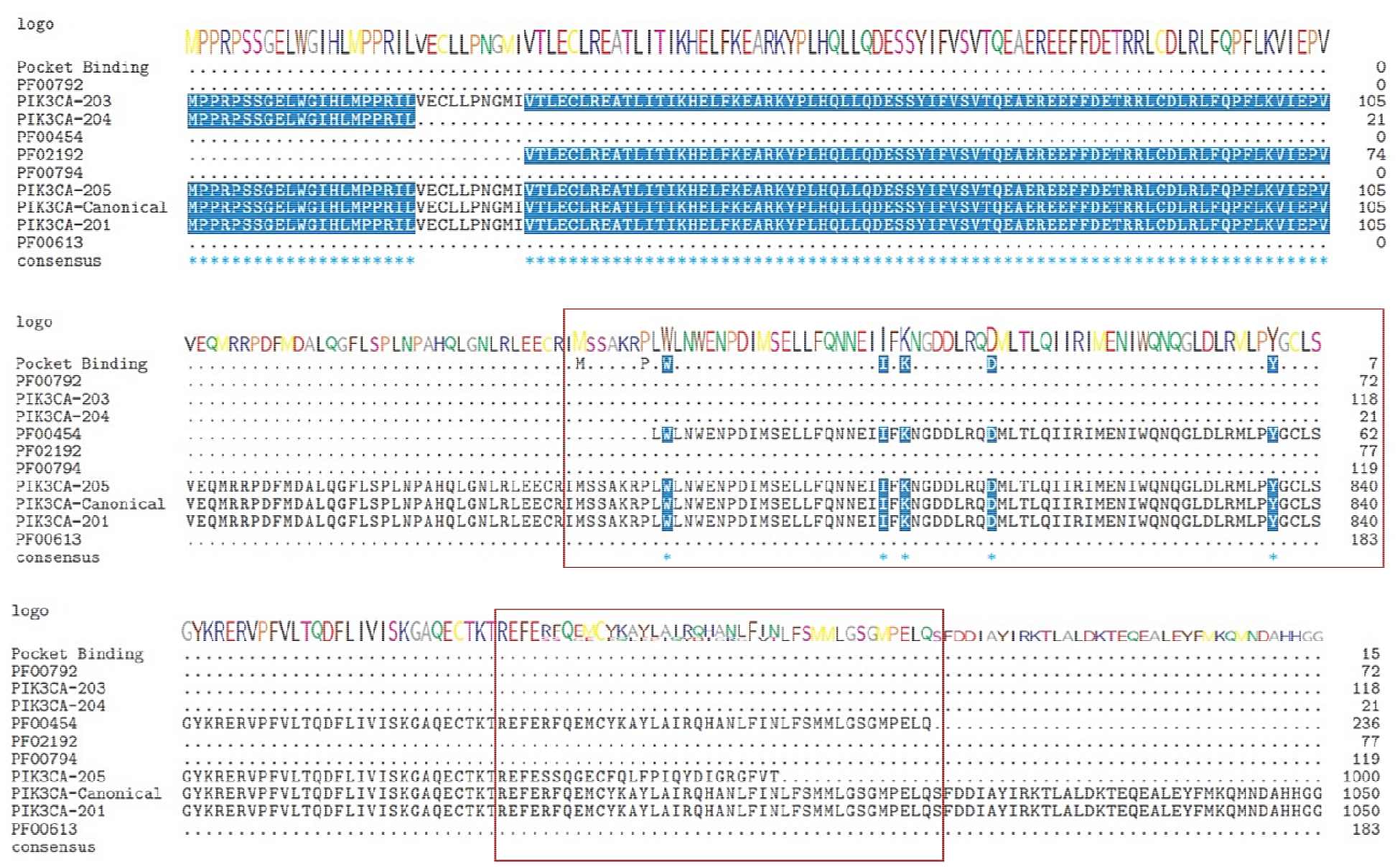
Sequence alignments of the predicted pocket binding residues of several PIK3CA protein isoforms. Using the Bioconductor software msa, Cluster Omega was used to align the binding residues with the isoform sequences. Predicted binding pocket residues, aligned Pfam domains, and PIK3CA-201, PIK3CA-203, PIK3CA-204, and PIK3CA-205 are shown from top to bottom. Each line included the consensus sequences’ sequence logo at the top. Residues in a sequence that coincide with the anticipated binding residues are shown by blue shading. The purple coloring suggests that this residue is conserved in about 50% of all sequences. Similar amino acids are shown by pink shading.

The KRAS gene has been a key target of cancer treatment discovery for decades since it is the most often mutated oncogene in human malignancies, notably in tumors of the pancreas, colon, and lung. However, despite these enormous efforts, cancers with KRAS mutations continue to be among the hardest to treat, in large part due to the emergence of treatment resistance brought on by the plasticity of tumor cells and/or the acquisition of additional mutations. According to the multiple sequence alignment of KRAS isoforms, the isoforms KRAS-203, 204 & 207 lack the binding pockets and are thus not predicted to be targets of drugs that treat the KRAS protein shown in Fig. 3. While the isoforms KRAS-201,202, 205, 210 and 214 have the same binding residues and are thus likely to be targeted by drugs. Further investigation revealed that KRAS-202,205,203, and 204 have variations with KRAS-201 on the C terminal. Our findings indicat that further effort is required to specifically target the KRAS isoforms.

**Fig. 3.**
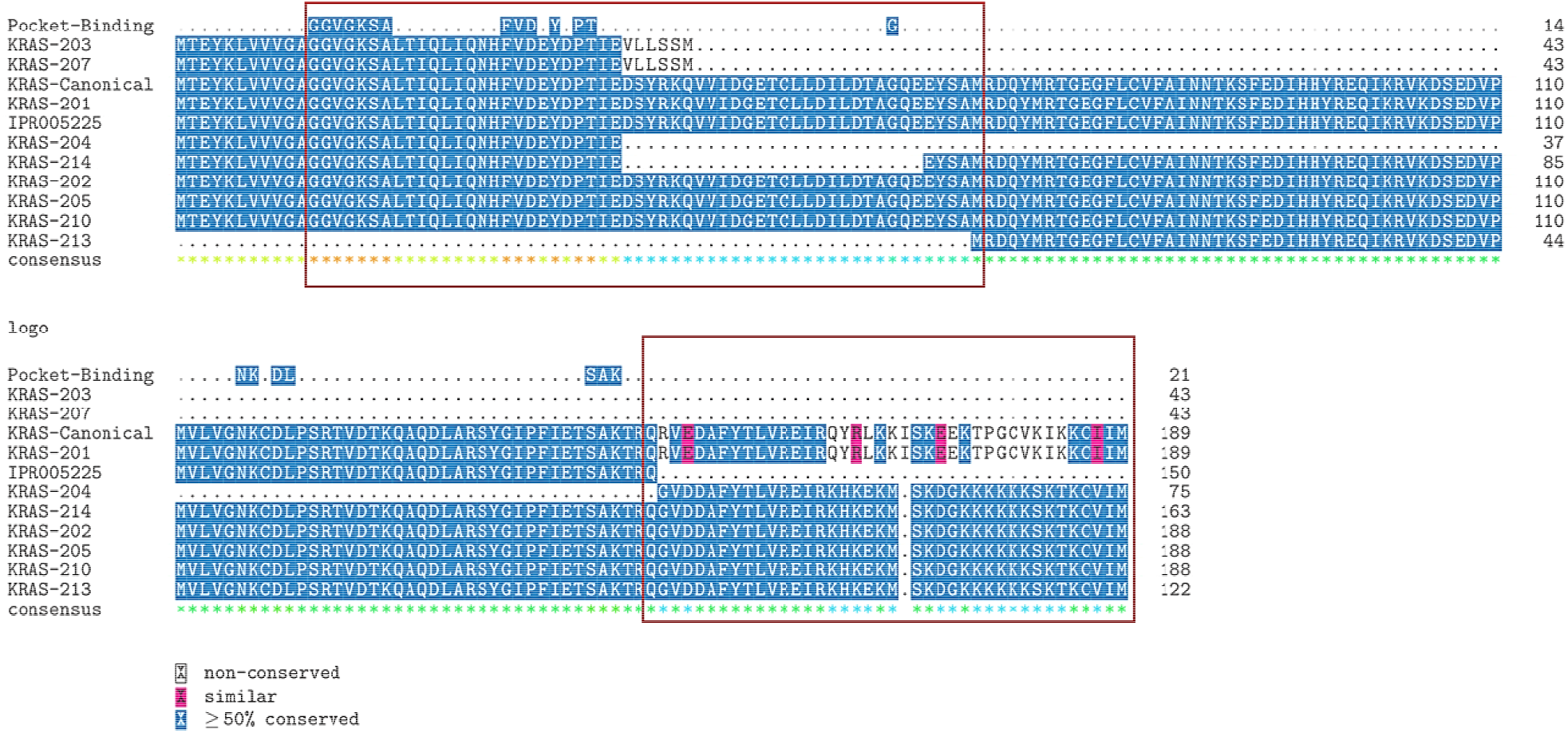
**A** Sequence alignments of predicted pocket binding residues on various KRAS protein isoforms. Using the Bioconductor software msa, Cluster Omega was used to align the binding residues with the isoform sequences. Predicted binding pocket residues, aligned Pfam domains and KRAS isoforms are shown from top to bottom. Each line included the consensus sequences’ sequence logo at the top. Residues in a sequence that coincide with the anticipated binding residues are shown by blue shading. The purple coloring suggests that this residue is conserved in about 50% of all sequences. Similar amino acids are shown by pink shading.

### 3.3 High Levels of Isoform Expression in Tumor Tissues

Using clinical information from UCSC Xena that is accessible through several projects (TCGA, GTEx and TARGET), we were able to determine the expression of protein isoforms. In TCGA samples of cervical cancer and breast cancer, we observed the expression of PIK3CA and KRAS isoforms shown in Fig. 4A. The expression of isoforms was nearly same in both cancer types. The isoform (PIK3CA-204/ENST00000477735.1) does not express in tumor and normal samples, and is thus ignored. The isoform (PIK3CA-203/ ENST00000468036.1) is highly expressed in the TCGA tumor samples, in contrast to the normal GTEx samples. While we previously found that isoform-203 does not have the predicted binding pocket but we observed that tumor cells express it. Thus, this should be included in future study to examines the on- and off-target effects of drugs.

**Fig. 4.**
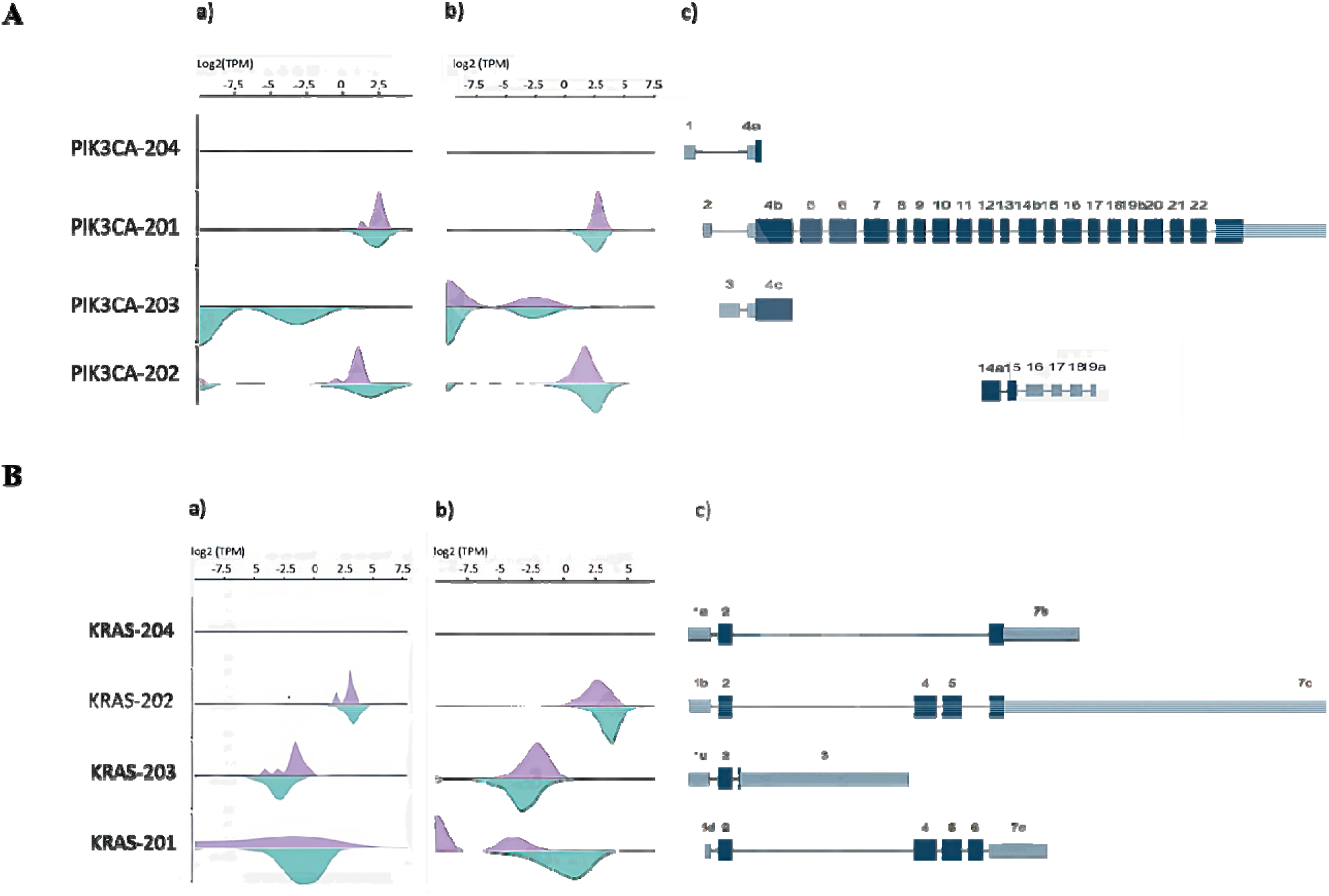
**A** PIK3CA isoform expression and exon structure. Green density represents log2(TPM) from GTEx normal samples, whereas purple density represents those from a) TCGA Cervical Cancer samples and b) TCGA Breast Cancer samples. Density plots and c) the exon structure plot both follow the same sequence. **B** KRAS isoform expression and exon structure. Four isoforms are related (from top to bottom). Green density represents those from GTEx normal samples, whereas purple density means a) TCGA Cervical Cancer samples and b) TCGA Breast Cancer Samples. Density plots and c) the exon structure plot both follow the same sequence. Every plot is generated using the UCSC Xena browser (37).

Using transcriptome expression data from the TCGA repository, it was possible to compare the expression of KRAS isoforms (KRAS-202/ENST00000311936.7, KRAS-203/ENST00000556131.1, and KRAS-204/ENST00000557334.5) in cervical and breast samples Fig. 4B. In comparison to normal samples, tumor samples were shown to have higher levels of KRAS-203 expression. Sequence analysis of FBXW7, ERBB3 & SMAD4 are shown in supplementary file 3. Future studies analyzing the on- and off-target effects of drugs should consider these isoforms as these are expressed in tumors

### 3.4 Drugs Interaction on Structural Level

Even though we have shown changes in binding pockets across isoforms at the sequence level, structural-level research is the only way to gain more solid proof that the drugs bind to their targets’ isoforms in distinct ways. We have studied the KRAS gene, which has seven distinct isoforms, together with known drugs that target them in to understand how a certain drug molecule interacts with several isoforms of a protein.

The 3D structures of each isoform were predicted using various databases. The best predicted structures were projected to have ERRAT scores greater than 94. While structures with poor ERRAT values were further improved.

Then, using Pyrex, we conducted docking analysis while taking into account a selection of drugs that have been identified to target this disease protein target. After analyzing the docked positions, we observed that although some drugs bind similarly to isoforms, while others bind extremely differently. For instance, Isoforms KRAS-203, 204 & 207 showed low binding affinity with the FDA Approved drugs (Table 2). It supports our previous findings that these isoforms have very small sequences and do not have the predicted binding pocket. While all other isoforms of KRAS (KRAS-201, 202, 205, 210, 213 & 214) have high binding affinities. AZD-4785 had good scores for KRAS-201, 202, 205, and 214. These six isoforms of the protein had strong binding affinity against Trametinib, although KRAS-202 had low binding affinity. With ridoforolimus, all isoforms had the good binding affinities. While the remaining drugs likewise shown good binding affinities with these isoforms, certain isoforms displayed lower affinities than others.

**Table 2.** Binding Affinity Values of the KRAS-Canonical protein and its isoforms.

| Drugs | KRAS-Canonical | KRAS-201 | KRAS-202 | KRAS-205 | KRAS-214 | KRAS-213 | KRAS-210 | KRAS-203 | KRAS-204 |
| --- | --- | --- | --- | --- | --- | --- | --- | --- | --- |
| **AZD-4785** | -6.9 | -7.3 | -7.2 | -7 | -7.2 | -6.4 | -7.9 | -5.7 | -4 |
| **AZD-8835** | -8.3 | -8.2 | -8.2 | -8.7 | -7.9 | -7.7 | -8.5 | -6.3 | -4.9 |
| **CC-223** | -7.2 | -7.7 | -7.5 | -7.5 | -7.7 | -7.3 | -7.7 | -6.1 | -4.5 |
| **PD-0325901** | -7.7 | -7.3 | -6.9 | -6.9 | -7.4 | -6.3 | -6.8 | -5.4 | -4.3 |
| **Ridaforolimus** | -10.1 | -10.2 | -9.7 | -9.6 | -10.6 | -10.7 | -9.6 | -8.8 | -5.3 |
| **Selumetinib** | -7.2 | -7.2 | -7.6 | -7.7 | -7.1 | -6.6 | -7.4 | -5.9 | -4.8 |
| **Trametinib** | -8.2 | -9 | -7.9 | -9.6 | -9 | -8.6 | -8.9 | -6.9 | -5.5 |

In case of PIK3CA, the isoforms PIK3CA-203 & 204 showed low binding affinity with approved FDA Drugs as these isoforms have short sequences and did not have predicted binding pocket (Table 3). While the isoforms PIK3CA-201 & 205 showed the best binding affinity with drugs. Temsirolimus showed good binding affinity with all isoforms

**Table 3.** Binding affinity values of the PIK3CA-Canonical, PIK3CA-201,205,203 & 204.

| **Drugs** | **PIK3CA-Canonical** | **PIK3CA-201** | **PIK3CA-205** | **PIK3CA-203** | **PIK3CA-204** |
| --- | --- | --- | --- | --- | --- |
| **CC-223** | -8.6 | -8.3 | -8 | -6.4 | -6.4 |
| **ALPELISIB** | -8.9 | -8.8 | -9.5 | -7.5 | -7.7 |
| **BUPARLISIB** | -8.6 | -8.2 | -8.1 | -6.2 | -6.4 |
| **CAPIVASERTIB** | -9.5 | -9.6 | -8.9 | -6.6 | -6.6 |
| **INK-1117** | -9 | -9 | -9 | -6.8 | -6.8 |
| **SERABELISIB** | -8.9 | -9.1 | -9 | -6.8 | -6.8 |
| TASELISIB | -9.7 | -9.7 | -8.2 | -7.2 | -6.9 |
| TEMSIROLIMUS | -9.5 | -11.6 | -10.2 | -9 | -9 |
| TRASTUZUMAB | -10.5 | -9.6 | -9.6 | -6.8 | -7.7 |

To explain how different pocket sizes, shapes, and electrostatic potential surfaces may create th illusion like the binding mode is different even when the scores are the same in some instances. Here, we examined Temsirolimus binding mode in all fours isoforms and discovered that while the binding scores are close, the binding patterns vary greatly shown in (Fig. 5). Molecular docking results of FBXW7, ERBB3 & SMAD4 are shown in supplementary file 4. These results led us to the hypothesis as, despite the identicality of the ligand binding residues, the binding pocket structures change in size, form, and dynamic properties, resulting in different binding patterns for a single drug in several isoforms with various binding affinity values.

**Fig. 5.**
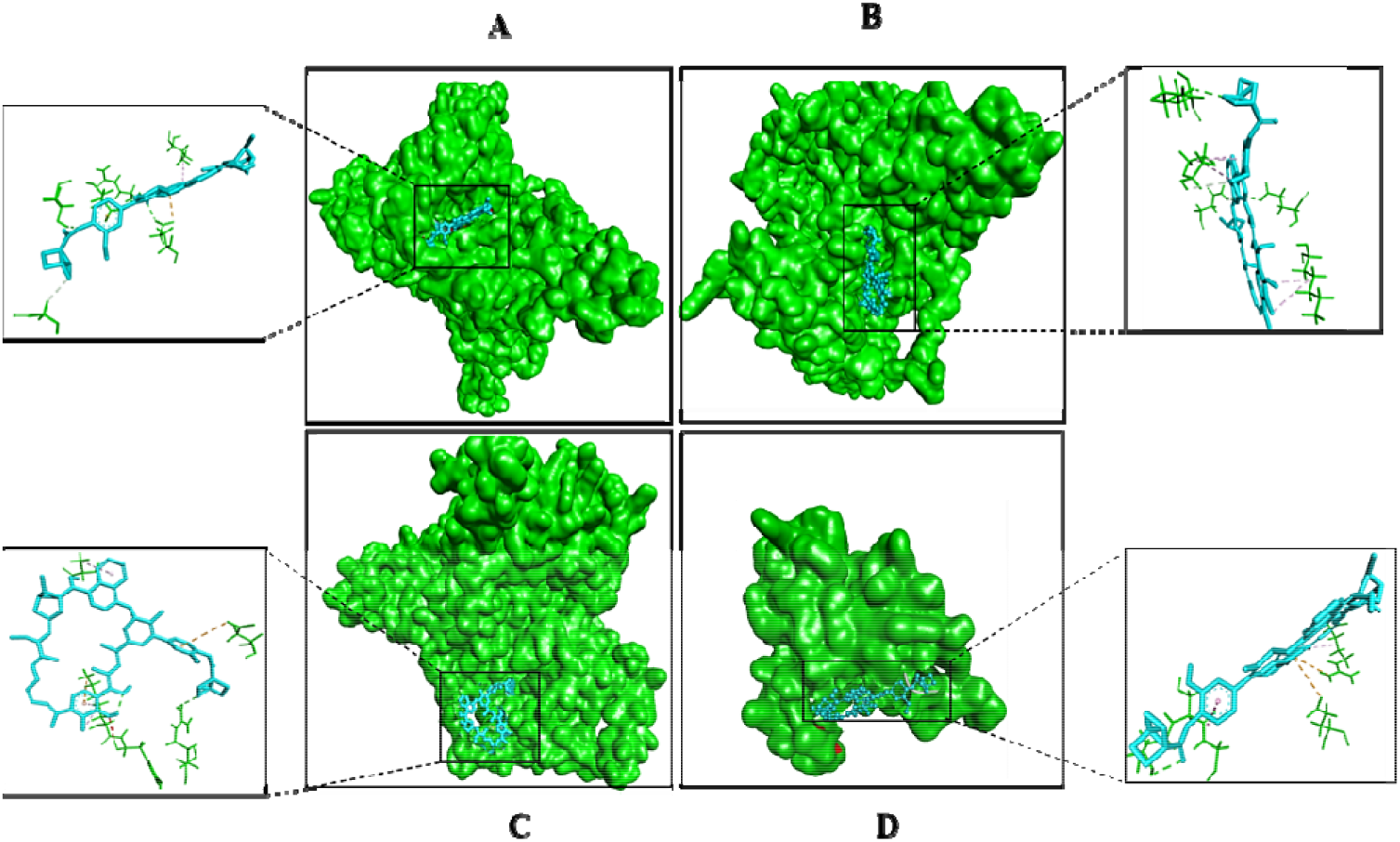
shows the ligand binding pocket of PIK3CA isoforms A) Canonical Protein B) PIK3CA-201 C) PIK3CA-205 and D) PIK3CA-202 with the drug Temsirolimus.

The interaction analysis of the target proteins isoforms was checked to see what kinds and how many interactions there were between the docked tesmilorous and the PIK3CA isoforms. When a complex has a significant number of hydrogen bonds together with a small number of salt bridges, hydrophobic contacts, and pi-pi interactions, it is said to be strong. To determine how many interactions each molecule generated, we tested each docked drug differently Fig. 6. According to the interaction study, complexes with strong binding affinities were those that produced the most hydrogen bonds (Table 4).

**Table 4.** Shows the Hydrogen and Hydrophobic interactions of docked isoforms with drugs.

| Protein | Hydrogen Interactions | Hydrophobic Interactions |
| --- | --- | --- |
| PIK3CA-Canonical | GLU, ASN, ASP, ASP, TYR | THR |
| PIK3CA-201 | ASP, ASN, LYS, SER | LYS, ASP, ASN, PRO, GLN |
| PIK3CA-205 | ARG, ASP, ASP, LYS, PHE | GLU, TYR, LYS |
| PIK3CA-203 | SER,THR | ARG, GLU |

**Fig. 6.**
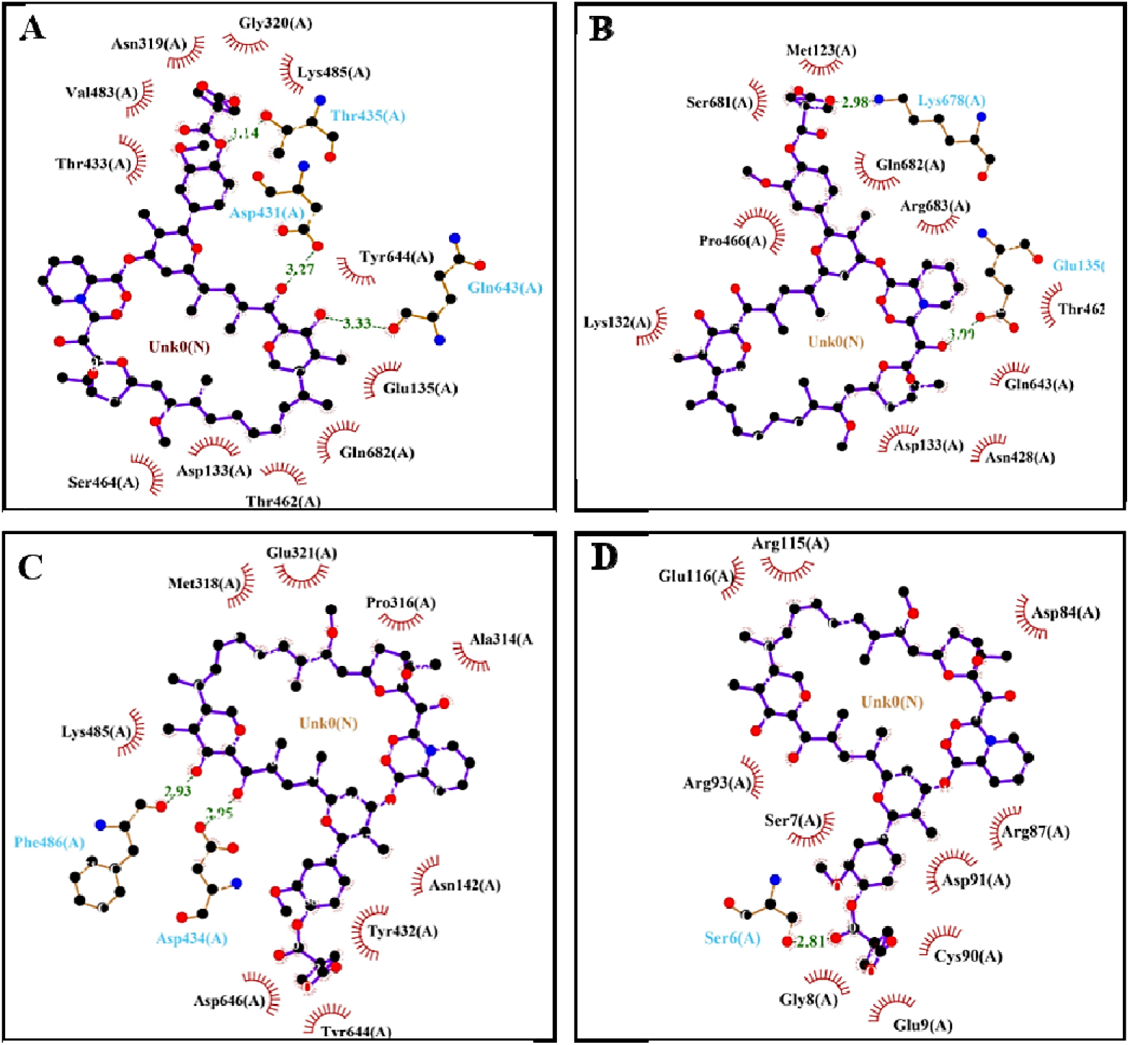
Ligplot analysis of interactions between PIK3CA isoforms and Temsirolimus. Hydrophobic interactions between amino acid residues are shown by red arcs, whereas hydrogen bonds are represented by green dashed lines with specified bond lengths.

PIK3CA-Canonical and isoforms 201 & 205 were shown to have strong interactions while the docked complex of PIK3CA-203 was found to have weak interactions.

## 4. Discussion

Despite the fact that current target prediction methods have shown the accuracy of genomic, chemical, and pharmacological data in drug target interaction prediction, those methods frequently only concentrate on the canonical isoforms while disregarding the on-or even off target isoform-level interactions that are linked to the chemical’s action (38). Previous research has related cancer-specific aberrant splicing to drug resistance mechanisms. However, little is known about the drug’s therapeutic impact on the specified tissue and its side effects on other tissues. Protein isoforms produced by alternative splicing can express at different levels and exhibit various, perhaps conflicting, activities in various tissues and/or organs (39, 40), We postulated in this study that various protein isoforms formed by alternative splicing might develop into candidates for drug interactions that are off-target or non-target because of the presence or lack of target binding sequence in different alternative splicing of genes specifically involved in cervical cancer. Our findings show that most small molecule therapeutic targets have a variety of protein isoforms. As a result, It’s therefore feasible which the most of pharmacological targeting genes’ protein isoforms have functional differences and show isoform-level changes in its interactions with the drug.

As we revealed that KRAS-203 is highly expressed in tumour samples, sequence alignment and data analysis of the gene expression patterns in the TCGA and GTEx datasets uncovered significant data, like medicines that skip alternative isoforms that also expressed in cancer but perhaps are not targeted, while the drugs which might possibly aim alternative isoforms that are variously expressed across many normal tissues, and those are involved in the process of cancer development. Furthermore, the same medication’s ability to bind to several structurally related isoforms with various affinities was verified by drug docking study and structural analysis of an example KRAS and PIK3CA protein. These findings basically two processes in which both possibly lead to far-off impacts, which could result in drug resistance.

In comparison to the canonical isoform, we observed low expression of KRAS isoforms in TCGA samples. We observed via structural docking research that various medicines can interact with all isoforms in various ways. It is still unknown whether the secondary isoforms behave similarly to or differently from the downregulated primary isoform, carcinogenic, as well as overexpressed. On the other hand, the different isoforms, with the exception of KRAS-204, which was not expressed in normal or tumor samples, showed variable and greater expression in healthy tissue than in tumor tissues. These isoforms can act as tumor suppressors or regulator, counteracting the carcinogenic isoform’s function. Immediate inhibition of these isoforms may be undesirable under such conditions. Despite the fact that the precise roles of these isoforms are yet unknown, it’s feasible because separating sites from non-targets at the splice level is a crucial step in early stages of drug discovery investigations.

Due to restrictions on the availability of data, we were challenged to have several limitations in our current study. The first challenge is the lack of mappings of isoforms between the public online database and older studies. For examples, there is frequently a difference in the exon numbers reported by these two sources. Public databases like Ensemble did not contain many of the isoforms that had previously been described in literature. This makes it extremely challenging to annotate these isoforms structurally and functionally. Therefore, the major aspects of our study are the overexpression of isoforms that are more advantageous for the development of cancer should be suppressed, and the main aims for suppression should be those isoforms that are upregulated in cancer. This is obviously a restriction because these two hypotheses might be incorrect, but as of right now, we don’t have any better methods for evaluating the roles of these unidentified isoforms. Furthermore, if there is inclusion of actual protein-level expression of these isoforms will strengthen the claim. As far as we are known, there is currently no comprehensive database that includes the expression of all protein isoforms on a complete proteome scale. In our opinion, the importance of comprehending pharmacological targets at the isoform level should be emphasized even more. However, our results add to those of a recent study that identified means mRNA expression across tissues and variance of expression across tissues as the two key characteristics that separate effective medications from ineffective ones (41).

## 5. Conclusion

We expect that our findings will encourage more future investigation into the possibility of isoform-level medication design. Enough structural and functional knowledge of these isoforms is necessary to accomplish this aim. Strongly identifying additional cancer biomarkers at the isoform level and connecting them to treatment sensitivity using computational methods would be a crucial next step. If isoform-level drug design is required, accurate structural modelling and prediction of these isoforms are particularly crucial because no database presently has such information about the structure in a well-annotated way. Additionally, various databases should continue to combine isoform-level information and analysis with earlier works of literature and ensure that they are in line, particularly with regard to the functional analyses of less common isoforms.

## Acknowledgments

I would like to acknowledge the substantial contribution of Muhammad Sajid in providing valuable resources and overseeing the supervision of this project. His guidance and expertise were instrumental in shaping the direction of our work.

