## Supplementary File 2 for "In Silico Analysis of Drug Off-Target Effects on Diverse Isoforms of Cervical Cancer for Enhanced Therapeutic Strategies"

```
Pocket Binding     .........................................................................................................    0
USP13 PIK3CA 499aa MQRRGALFGMPGGSGGRKMAAGDIGELLVPHMPTIRVPRSGDRVYKNECAFSYDSPNSEGGLYVCMNTFLAFGREHVERHFRKTGQSVYMHLKRHVREKVRGASG  105
USP13 PIK3CA 434aa .................................................................MNTFLAFGREHVERHFRKTGQSVYMHLKRHVREKVRGASG   40
consensus                                                                           ****************************************

Pocket Binding     ................................................................................................M.....P.W    3
USP13 PIK3CA 499aa GALPKRRNSKIFLDLDTDDDLNSDDYEYEDEAKLVIFPDHYEIALPNIEELPALVQMKFLVEQMRRPDFMDALQGFLSPLNPAHQLGNLRLEECRIMSSAKRPLW  210
USP13 PIK3CA 434aa GALPKRRNSKIFLDLDTDDDLNSDDYEYEDEAKLVIFPDHYEIALPNIEELPALVQMKFLVEQMRRPDFMDALQGFLSPLNPAHQLGNLRLEECRIMSSAKRPLW  145
consensus          ************************************************************************************************!*****!*!

Pocket Binding     ...................I.K.......D.........................Y...........IE.V....T.............................   11
USP13 PIK3CA 499aa LNWENPDIMSELLFQNNEIIFKNGDDLRQDMLTLQIIRIMENIWQNQGLDLRMLPYGCLSIGDCVGLIEVVRNSHTIMQIQCKGGLKGALQFNSHTLHQWLKDKN  315
USP13 PIK3CA 434aa LNWENPDIMSELLFQNNEIIFKNGDDLRQDMLTLQIIRIMENIWQNQGLDLRMLPYGCLSIGDCVGLIEVVRNSHTIMQIQCKGGLKGALQFNSHTLHQWLKDKN  250
consensus          *******************!*!*******!*************************!***********!!*!****!*****************************

Pocket Binding     ....................................M.......F.ID.........................................................   15
USP13 PIK3CA 499aa KGEIYDAAIDLFTRSCAGYCVATFILGIGDRHNSNIMVKDDGQLFHIDFGHFLDHKKKKFGYKRERVPFVLTQDFLIVISKGAQECTKTREFERFQEMCYKAYLA  420
USP13 PIK3CA 434aa KGEIYDAAIDLFTRSCAGYCVATFILGIGDRHNSNIMVKDDGQLFHIDFGHFLDHKKKKFGYKRERVPFVLTQDFLIVISKGAQECTKTREFERFQEMCYKAYLA  355
consensus          ************************************!*******!*!!*********************************************************

Pocket Binding     ..............................   15
USP13 PIK3CA 499aa IRQHANLFINLFSMMLGSGMPELQSFDDIA  450
USP13 PIK3CA 434aa IRQHA.........................  360
consensus          *****

X  non-conserved
X  ≥50% conserved
```
