## Supplementary File 3 for "In Silico Analysis of Drug Off-Target Effects on Diverse Isoforms of Cervical Cancer for Enhanced Therapeutic Strategies"

@
@
@

@
@

logo 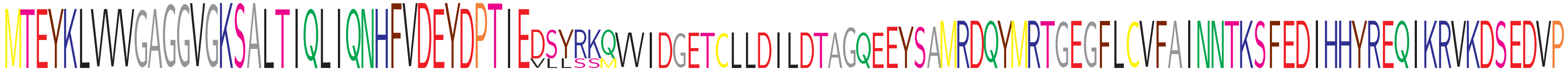

```
Pocket-Binding  ...........GGVGKSA.........FVD.Y.PT........................G..................................................   14
KRAS-203        MTEYKLVVVGAGGVGKSALTIQLIQNHFVDEYDPTIEVLLSSM...................................................................   43
KRAS-207        MTEYKLVVVGAGGVGKSALTIQLIQNHFVDEYDPTIEVLLSSM...................................................................   43
KRAS-Canonical  MTEYKLVVVGAGGVGKSALTIQLIQNHFVDEYDPTIEDSYRKQVVIDGETCLLDILDTAGQEEYSAMRDQYMRTGEGFLCVFAINNTKSFEDIHHYREQIKRVKDSEDVP  110
KRAS-201        MTEYKLVVVGAGGVGKSALTIQLIQNHFVDEYDPTIEDSYRKQVVIDGETCLLDILDTAGQEEYSAMRDQYMRTGEGFLCVFAINNTKSFEDIHHYREQIKRVKDSEDVP  110
IPR005225       MTEYKLVVVGAGGVGKSALTIQLIQNHFVDEYDPTIEDSYRKQVVIDGETCLLDILDTAGQEEYSAMRDQYMRTGEGFLCVFAINNTKSFEDIHHYREQIKRVKDSEDVP  110
KRAS-204        MTEYKLVVVGAGGVGKSALTIQLIQNHFVDEYDPTIE.........................................................................   37
KRAS-214        MTEYKLVVVGAGGVGKSALTIQLIQNHFVDEYDPTIE.........................EYSAMRDQYMRTGEGFLCVFAINNTKSFEDIHHYREQIKRVKDSEDVP   85
KRAS-202        MTEYKLVVVGAGGVGKSALTIQLIQNHFVDEYDPTIEDSYRKQVVIDGETCLLDILDTAGQEEYSAMRDQYMRTGEGFLCVFAINNTKSFEDIHHYREQIKRVKDSEDVP  110
KRAS-205        MTEYKLVVVGAGGVGKSALTIQLIQNHFVDEYDPTIEDSYRKQVVIDGETCLLDILDTAGQEEYSAMRDQYMRTGEGFLCVFAINNTKSFEDIHHYREQIKRVKDSEDVP  110
KRAS-210        MTEYKLVVVGAGGVGKSALTIQLIQNHFVDEYDPTIEDSYRKQVVIDGETCLLDILDTAGQEEYSAMRDQYMRTGEGFLCVFAINNTKSFEDIHHYREQIKRVKDSEDVP  110
KRAS-213        ..................................................................MRDQYMRTGEGFLCVFAINNTKSFEDIHHYREQIKRVKDSEDVP   44
consensus       **************************************************************************************************************
```

logo 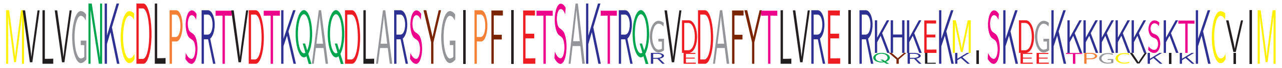

```
Pocket-Binding  .....NK.DL........................SAK..........................................   21
KRAS-203        ...............................................................................   43
KRAS-207        ...............................................................................   43
KRAS-Canonical  MVLVGNKCDLPSRTVDTKQAQDLARSYGIPFIETSAKTRQRVEDAFYTLVREIRQYRLKKISKEEKTPGCVKIKKCIIM  189
KRAS-201        MVLVGNKCDLPSRTVDTKQAQDLARSYGIPFIETSAKTRQRVEDAFYTLVREIRQYRLKKISKEEKTPGCVKIKKCIIM  189
IPR005225       MVLVGNKCDLPSRTVDTKQAQDLARSYGIPFIETSAKTRQ.......................................  150
KRAS-204        ........................................GVDDAFYTLVREIRKHKEKM.SKDGKKKKKKSKTKCVIM   75
KRAS-214        MVLVGNKCDLPSRTVDTKQAQDLARSYGIPFIETSAKTRQGVDDAFYTLVREIRKHKEKM.SKDGKKKKKKSKTKCVIM  163
KRAS-202        MVLVGNKCDLPSRTVDTKQAQDLARSYGIPFIETSAKTRQGVDDAFYTLVREIRKHKEKM.SKDGKKKKKKSKTKCVIM  188
KRAS-205        MVLVGNKCDLPSRTVDTKQAQDLARSYGIPFIETSAKTRQGVDDAFYTLVREIRKHKEKM.SKDGKKKKKKSKTKCVIM  188
KRAS-210        MVLVGNKCDLPSRTVDTKQAQDLARSYGIPFIETSAKTRQGVDDAFYTLVREIRKHKEKM.SKDGKKKKKKSKTKCVIM  188
KRAS-213        MVLVGNKCDLPSRTVDTKQAQDLARSYGIPFIETSAKTRQGVDDAFYTLVREIRKHKEKM.SKDGKKKKKKSKTKCVIM  122
consensus       ************************************************************ ******************
```

X non-conserved
X similar
X ≥50% conserved

```
PF00454             .........................................................................................................   0
PF02192             ...............................VTLECLREATLITIKHELFKEARKYPLHQLLQDESSYIFVSVTQEAEREEFFDETRRLCDLRLFQPFLKVIEPV  74
PF00794             .........................................................................................................   0
PIK3CA-205          MPPRPSSGELWGIHLMPPRILVECLLPNGMIVTLECLREATLITIKHELFKEARKYPLHQLLQDESSYIFVSVTQEAEREEFFDETRRLCDLRLFQPFLKVIEPV 105
PIK3CA-Canonical    MPPRPSSGELWGIHLMPPRILVECLLPNGMIVTLECLREATLITIKHELFKEARKYPLHQLLQDESSYIFVSVTQEAEREEFFDETRRLCDLRLFQPFLKVIEPV 105
PIK3CA-201          MPPRPSSGELWGIHLMPPRILVECLLPNGMIVTLECLREATLITIKHELFKEARKYPLHQLLQDESSYIFVSVTQEAEREEFFDETRRLCDLRLFQPFLKVIEPV 105
PIK3CA-204          .........................................................................................................  21
PF00454             .........................................................................................................   0
PF02192             GNR......................................................................................................  77
PF00794             ....................................................................SPELPKHIYNKLDKGQIIVVIWVIVSPNNDKQKYTLK  37
PIK3CA-205          GNREEKILNREIGFAIGMPVCEFDMVKDPEVQDFRRNILNVCKEAVDLRDLNSPHSRAMYVYPPNVESSPELPKHIYNKLDKGQIIVVIWVIVSPNNDKQKYTLK 210
PIK3CA-Canonical    GNREEKILNREIGFAIGMPVCEFDMVKDPEVQDFRRNILNVCKEAVDLRDLNSPHSRAMYVYPPNVESSPELPKHIYNKLDKGQIIVVIWVIVSPNNDKQKYTLK 210
PIK3CA-201          GNREEKILNREIGFAIGMPVCEFDMVKDPEVQDFRRNILNVCKEAVDLRDLNSPHSRAMYVYPPNVESSPELPKHIYNKLDKGQIIVVIWVIVSPNNDKQKYTLK 210
PIK3CA-204          .........................................................................................................  21
PF00454             .........................................................................................................   0
PF02192             .........................................................................................................  77
PF00794             INHDCVPEQVIAEAIRKKTRSMLLSSEQLKLCVLEYQGKYILKVCGCDEYFLEKYPLSQYKYIRSCIMLGRMPNLMLMAKES....................... 119
PIK3CA-205          INHDCVPEQVIAEAIRKKTRSMLLSSEQLKLCVLEYQGKYILKVCGCDEYFLEKYPLSQYKYIRSCIMLGRMPNLMLMAKESLYSQLPMDCFTMPSYSRRISTAT 315
PIK3CA-Canonical    INHDCVPEQVIAEAIRKKTRSMLLSSEQLKLCVLEYQGKYILKVCGCDEYFLEKYPLSQYKYIRSCIMLGRMPNLMLMAKESLYSQLPMDCFTMPSYSRRISTAT 315
PIK3CA-201          INHDCVPEQVIAEAIRKKTRSMLLSSEQLKLCVLEYQGKYILKVCGCDEYFLEKYPLSQYKYIRSCIMLGRMPNLMLMAKESLYSQLPMDCFTMPSYSRRISTAT 315
PF00613             .........................................................................................................   0
consensus

Pocket Binding      .........................................................................................................   0
PF00792             ................................................................................................DKIYVRTGI   9
PIK3CA-203          ......................................................................................................... 118
PIK3CA-204          .........................................................................................................  21
PF00454             .........................................................................................................   0
PF02192             .........................................................................................................  77
PF00794             ......................................................................................................... 119
PIK3CA-205          PYMNGETSTKSLWVINSALRIKILCATYVNVNIRDIDKIYVRTGIYHGGEPLCDNVNTQRVPCSNPRWNEWLNYDIYIPDLPRAARLCLSICSVKGRKGAKEEHC 420
PIK3CA-Canonical    PYMNGETSTKSLWVINSALRIKILCATYVNVNIRDIDKIYVRTGIYHGGEPLCDNVNTQRVPCSNPRWNEWLNYDIYIPDLPRAARLCLSICSVKGRKGAKEEHC 420
PIK3CA-201          PYMNGETSTKSLWVINSALRIKILCATYVNVNIRDIDKIYVRTGIYHGGEPLCDNVNTQRVPCSNPRWNEWLNYDIYIPDLPRAARLCLSICSVKGRKGAKEEHC 420
PF00613             .........................................................................................................   0
consensus
```

```
Pocket Binding    .........................................................................................................    0
PF00792           PLAWGNINLFDYTDTLVSGKMALNLWPVPHGLEDLLNPIGVTGSNPNKETPCLELEFDWFSSV..........................................   72
PIK3CA-203        .........................................................................................................  118
PIK3CA-204        .........................................................................................................   21
PF00454           .........................................................................................................    0
PF02192           .........................................................................................................   77
PF00794           .........................................................................................................  119
PIK3CA-205        PLAWGNINLFDYTDTLVSGKMALNLWPVPHGLEDLLNPIGVTGSNPNKETPCLELEFDWFSSVVKFPDMSVIEEHANWSVSREAGFSYSHAGLSNRLARDNELRE  525
PIK3CA-Canonical  PLAWGNINLFDYTDTLVSGKMALNLWPVPHGLEDLLNPIGVTGSNPNKETPCLELEFDWFSSVVKFPDMSVIEEHANWSVSREAGFSYSHAGLSNRLARDNELRE  525
PIK3CA-201        PLAWGNINLFDYTDTLVSGKMALNLWPVPHGLEDLLNPIGVTGSNPNKETPCLELEFDWFSSVVKFPDMSVIEEHANWSVSREAGFSYSHAGLSNRLARDNELRE  525
PF00613           .....................................................................................................ELRE    4
consensus

Pocket Binding    .........................................................................................................    0
PF00792           .........................................................................................................   72
PIK3CA-203        .........................................................................................................  118
PIK3CA-204        .........................................................................................................   21
PF00454           .........................................................................................................    0
PF02192           .........................................................................................................   77
PF00794           .........................................................................................................  119
PIK3CA-205        NDKEQLKAISTRDPLSEITEQEKDFLWSHRHYCVTIPEILPKLLLSVKWNSRDEVAQMYCLVKDWPPIKPEQAMELLDCNYPDPMVRGFAVRCLEKYLTDDKLSQ  630
PIK3CA-Canonical  NDKEQLKAISTRDPLSEITEQEKDFLWSHRHYCVTIPEILPKLLLSVKWNSRDEVAQMYCLVKDWPPIKPEQAMELLDCNYPDPMVRGFAVRCLEKYLTDDKLSQ  630
PIK3CA-201        NDKEQLKAISTRDPLSEITEQEKDFLWSHRHYCVTIPEILPKLLLSVKWNSRDEVAQMYCLVKDWPPIKPEQAMELLDCNYPDPMVRGFAVRCLEKYLTDDKLSQ  630
PF00613           NDKEQLKAISTRDPLSEITEQEKDFLWSHRHYCVTIPEILPKLLLSVKWNSRDEVAQMYCLVKDWPPIKPEQAMELLDCNYPDPMVRGFAVRCLEKYLTDDKLSQ  109
consensus

Pocket Binding    .........................................................................................................    0
PF00792           .........................................................................................................   72
PIK3CA-203        .........................................................................................................  118
PIK3CA-204        .........................................................................................................   21
PF00454           .........................................................................................................    0
PF02192           .........................................................................................................   77
PF00794           .........................................................................................................  119
PIK3CA-205        YLIQLVQVLKYEQYLDNLLVRFLLKKALTNQRIGHFFFWHLKSEMHNKTVSQRFGLLLESYCRACGMYLKHLNRQVEAMEKLINLTDILKQEKKDETQKVQMKFL  735
PIK3CA-Canonical  YLIQLVQVLKYEQYLDNLLVRFLLKKALTNQRIGHFFFWHLKSEMHNKTVSQRFGLLLESYCRACGMYLKHLNRQVEAMEKLINLTDILKQEKKDETQKVQMKFL  735
PIK3CA-201        YLIQLVQVLKYEQYLDNLLVRFLLKKALTNQRIGHFFFWHLKSEMHNKTVSQRFGLLLESYCRACGMYLKHLNRQVEAMEKLINLTDILKQEKKDETQKVQMKFL  735
PF00613           YLIQLVQVLKYEQYLDNLLVRFLLKKALTNQRIGHFFFWHLKSEMHNKTVSQRFGLLLESYCRACGMYLKHLNR...............................  183
consensus

Pocket Binding    ....................................M.....P.W...................I.K.......D.........................Y....    7
PF00792           .........................................................................................................   72
PIK3CA-203        .........................................................................................................  118
PIK3CA-204        .........................................................................................................   21
PF00454           ...........................................LWLNWENPDIMSELLFQNNEIIFKNGDDLRQDMLTLQIIRIMENIWQNQGLDLRMLPYGCLS   62
PF02192           .........................................................................................................   77
PF00794           .........................................................................................................  119
PIK3CA-205        VEQMRRPDFMDALQGFLSPLNPAHQLGNLRLEECRIMSSAKRPLWLNWENPDIMSELLFQNNEIIFKNGDDLRQDMLTLQIIRIMENIWQNQGLDLRMLPYGCLS  840
PIK3CA-Canonical  VEQMRRPDFMDALQGFLSPLNPAHQLGNLRLEECRIMSSAKRPLWLNWENPDIMSELLFQNNEIIFKNGDDLRQDMLTLQIIRIMENIWQNQGLDLRMLPYGCLS  840
PIK3CA-201        VEQMRRPDFMDALQGFLSPLNPAHQLGNLRLEECRIMSSAKRPLWLNWENPDIMSELLFQNNEIIFKNGDDLRQDMLTLQIIRIMENIWQNQGLDLRMLPYGCLS  840
PF00613           .........................................................................................................  183
consensus                                                     *                   * *       *                         *
```

```
Pocket Binding    .......IE.V....T.................................................................M.......F.ID............   15
PF00792           .........................................................................................................   72
PIK3CA-203        .........................................................................................................  118
PIK3CA-204        .........................................................................................................   21
PF00454           IGDCVGLIEVVRNSHTIMQIQCKGGLKGALQFNSHTLHQWLKDKNKGEIYDAAIDLFTRSCAGYCVATFILGIGDRHNSNIMVKDDGQLFHIDFGHFLDHKKKKF  167
PF02192           .........................................................................................................   77
PF00794           .........................................................................................................  119
PIK3CA-205        IGDCVGLIEVVRNSHTIMQIQCKGGLKGALQFNSHTLHQWLKDKNKGEIYDAAIDLFTRSCAGYCVATFILGIGDRHNSNIMVKDDGQLFHIDFGHFLDHKKKKF  945
PIK3CA-Canonical  IGDCVGLIEVVRNSHTIMQIQCKGGLKGALQFNSHTLHQWLKDKNKGEIYDAAIDLFTRSCAGYCVATFILGIGDRHNSNIMVKDDGQLFHIDFGHFLDHKKKKF  945
PIK3CA-201        IGDCVGLIEVVRNSHTIMQIQCKGGLKGALQFNSHTLHQWLKDKNKGEIYDAAIDLFTRSCAGYCVATFILGIGDRHNSNIMVKDDGQLFHIDFGHFLDHKKKKF  945
PF00613           .........................................................................................................  183
consensus                **  *    *                                                                 *       * **

Pocket Binding    .........................................................................................................   15
PF00792           .........................................................................................................   72
PIK3CA-203        .........................................................................................................  118
PIK3CA-204        .........................................................................................................   21
PF00454           GYKRERVPFVLTQDFLIVISKGAQECTKTREFERFQEMCYKAYLAIRQHANLFINLFSMMLGSGMPELQ....................................  236
PF02192           .........................................................................................................   77
PF00794           .........................................................................................................  119
PIK3CA-205        GYKRERVPFVLTQDFLIVISKGAQECTKTREFESSQGECFQLFPIQYDIGRGFVT.................................................. 1000
PIK3CA-Canonical  GYKRERVPFVLTQDFLIVISKGAQECTKTREFERFQEMCYKAYLAIRQHANLFINLFSMMLGSGMPELQSFDDIAYIRKTLALDKTEQEALEYFMKQMNDAHHGG 1050
PIK3CA-201        GYKRERVPFVLTQDFLIVISKGAQECTKTREFERFQEMCYKAYLAIRQHANLFINLFSMMLGSGMPELQSFDDIAYIRKTLALDKTEQEALEYFMKQMNDAHHGG 1050
PF00613           .........................................................................................................  183
consensus

Pocket Binding    ..................   15
PF00792           ..................   72
PIK3CA-203        ..................  118
PIK3CA-204        ..................   21
PF00454           ..................  236
PF02192           ..................   77
PF00794           ..................  119
PIK3CA-205        .................. 1000
PIK3CA-Canonical  WTTKMDWIFHTIKQHALN 1068
PIK3CA-201        WTTKMDWIFHTIKQHALN 1068
PF00613           ..................  183
consensus
```

X non-conserved

X ≥50% conserved

@
@
@

@
@

logo
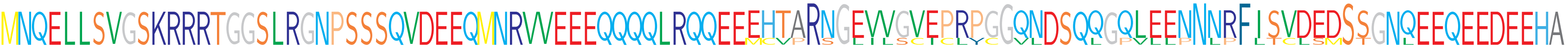

```
Pocket-Binding    .........................................................................................................   0
FBXW7-202         .........................................................................................................   0
FBXW7-203         ..................................................MCVPRSGLILSCICLYCGVL...L.PVLLPNLPFLTCLSMST..L..........  39
FBXW7-Canonical   MNQELLSVGSKRRRTGGSLRGNPSSSQVDEEQMNRVVEEEQQQQLRQQEEEHTARNGEVVGVEPRPGGQNDSQQGQLEENNNRFISVDEDSSGNQEEQEEDEEHA 105
FBXW7-201         MNQELLSVGSKRRRTGGSLRGNPSSSQVDEEQMNRVVEEEQQQQLRQQEEEHTARNGEVVGVEPRPGGQNDSQQGQLEENNNRFISVDEDSSGNQEEQEEDEEHA 105
FBXW7-204         MNQELLSVGSKRRRTGGSLRGNPSSSQVDEEQMNRVVEEEQQQQLRQQEEEHTARNGEVVGVEPRPGGQNDSQQGQLEENNNRFISVDEDSSGNQEEQEEDEEHA 105
FBXW7-206         MNQELLSVGSKRRRTGGSLRGNPSSSQVDEEQMNRVVEEEQQQQLRQQEEEHTARNGEVVGVEPRPGGQNDSQQGQLEENNNRFISVDEDSSGNQEEQEEDEEHA 105
IPR001810         .........................................................................................................   0
PF12937           .........................................................................................................   0
consensus         *********************************************************************************************************
```

logo
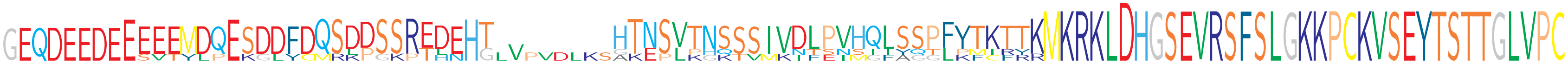

```
Pocket-Binding    ..................................V........................................D.............................   2
FBXW7-202         .....................MSKPGKPTLNHGLVPVDLKSAKEPLPHQTVMKIFSISIIAQGLPFCRRRMKRKLDHGSEVRSFSLGKKPCKVSEYTSTTGLVPC  84
FBXW7-203         ........ESVTYLPEKGLYCQRLPSSRTH..G........GTESLKGK....NTENMGFYGTLKMIFYKMKRKLDHGSEVRSFSLGKKPCKVSEYTSTTGLVPC 122
FBXW7-Canonical   GEQDEEDEEEEEMDQESDDFDQSDDSSREDEHT........HTNSVTNSSSIVDLPVHQLSSPFYTKTTKMKRKLDHGSEVRSFSLGKKPCKVSEYTSTTGLVPC 202
FBXW7-201         GEQDEEDEEEEEMDQESDDFDQSDDSSREDEHT........HTNSVTNSSSIVDLPVHQLSSPFYTKTTKMKRKLDHGSEVRSFSLGKKPCKVSEYTSTTGLVPC 202
FBXW7-204         GEQDEEDEEEEEMDQESDDFDQSDDSSREDEHT........HTNSVTNSSSIVDLPVHQLSSPFYTKTTKMKRKLDHGSEVRSFSLGKKPCKVSEYTSTTGLVPC 202
FBXW7-206         GEQDEEDEEEEEMDQESDDFDQSDDSSREDEHT........HTNSVTNSSSIVDLPVHQLSSPFYTKTTKMKRKLDHGSEVRSFSLGKKPCKVSEYTSTTGLVPC 202
IPR001810         .........................................................................................................   0
PF12937           .........................................................................................................   0
consensus         *********************************        ****************************************************************
```

logo
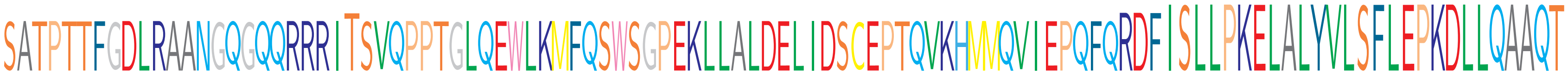

```
Pocket-Binding    .......................T.................................................................................   3
FBXW7-202         SATPTTFGDLRAANGQGQQRRRITSVQPPTGLQEWLKMFQSWSGPEKLLALDELIDSCEPTQVKHMMQVIEPQFQRDFISLLPKELALYVLSFLEPKDLLQAAQT 189
FBXW7-203         SATPTTFGDLRAANGQGQQRRRITSVQPPTGLQEWLKMFQSWSGPEKLLALDELIDSCEPTQVKHMMQVIEPQFQRDFISLLPKELALYVLSFLEPKDLLQAAQT 227
FBXW7-Canonical   SATPTTFGDLRAANGQGQQRRRITSVQPPTGLQEWLKMFQSWSGPEKLLALDELIDSCEPTQVKHMMQVIEPQFQRDFISLLPKELALYVLSFLEPKDLLQAAQT 307
FBXW7-201         SATPTTFGDLRAANGQGQQRRRITSVQPPTGLQEWLKMFQSWSGPEKLLALDELIDSCEPTQVKHMMQVIEPQFQRDFISLLPKELALYVLSFLEPKDLLQAAQT 307
FBXW7-204         SATPTTFGDLRAANGQGQQRRRITSVQPPTGLQEWLKMFQSWSGPEKLLALDELIDSCEPTQVKHMMQVIEPQFQRDFISLLPKELALYVLSFLEPKDLLQAAQT 307
FBXW7-206         SATPTTFGDLRAANGQGQQRRRITSVQPPTGLQEWLKMFQSWSGPEKLLALDELIDSCEPTQVKHMMQVIEPQFQRDFISLLPKELALYVLSFLEPKDLLQAAQT 307
IPR001810         ...........................................................................RDFISLLPKELALYVLSFLEPKDLLQAAQT  30
PF12937           ..............................................................................ISLLPKELALYVLSFLEPKDLLQAAQT  27
consensus         *********************************************************************************************************
```

```
logo               CRYWRILAEDNLLWREKCKEEGIDEPLHIKRRKVIKPGFIHSPWKSAYIRQHRIDTNWRRGELKSPKVLKGHDDHVITCLQFCGNRIVSGSDDNTLKVWSAVTGK
Pocket-Binding     .........................................................................................................   3
FBXW7-202          CRYWRILAEDNLLWREKCKEEGIDEPLHIKRRKVIKPGFIHSPWKSAYIRQHRIDTNWRRGELKSPKVLKGHDDHVITCLQFCGNRIVSGSDDNTLKVWSAVTGK 294
FBXW7-203          CRYWRILAEDNLLWREKCKEEGIDEPLHIKRRKVIKPGFIHSPWKSAYIRQHRIDTNWRRGELKSPKVLKGHDDHVITCLQFCGNRIVSGSDDNTLKVWSAVTGK 332
FBXW7-Canonical    CRYWRILAEDNLLWREKCKEEGIDEPLHIKRRKVIKPGFIHSPWKSAYIRQHRIDTNWRRGELKSPKVLKGHDDHVITCLQFCGNRIVSGSDDNTLKVWSAVTGK 412
FBXW7-201          CRYWRILAEDNLLWREKCKEEGIDEPLHIKRRKVIKPGFIHSPWKSAYIRQHRIDTNWRRGELKSPKVLKGHDDHVITCLQFCGNRIVSGSDDNTLKVWSAVTGK 412
FBXW7-204          CRYWRILAEDNLLWREKCKEEGIDEPLHIKRRKVIKPGFIHSPWKSAYIRQHRIDTNWRRGELKSPKVLKGHDDHVITCLQFCGNRIVSGSDDNTLKVWSAVTGK 412
FBXW7-206          CRYWRILAEDNLLWREKCKEEGIDEPLHIKRRKVIKPGFIHSPWKSAYIRQHRIDTNWRRGELKSPKVLKGHDDHVITCLQFCGNRIVSGSDDNTLKVWSAVTGK 412
IPR001810          CRYWRILAEDNLLWREKC.......................................................................................  48
PF12937            CRYWRILAEDNLLWREKC.......................................................................................  45
consensus          *********************************************************************************************************

logo               CLRTLVGHTGGVWSSQMRDNIIISGSTDRTLKVWNAETGECIHTLYGHTSTVRCMHLHEKRVVSGSRDATLRVWDIETGQCLHVLMGHVAAVRCVQYDGRRVVSG
Pocket-Binding     ....................................................R.............R.........................R............   6
FBXW7-202          CLRTLVGHTGGVWSSQMRDNIIISGSTDRTLKVWNAETGECIHTLYGHTSTVRCMHLHEKRVVSGSRDATLRVWDIETGQCLHVLMGHVAAVRCVQYDGRRVVSG 399
FBXW7-203          CLRTLVGHTGGVWSSQMRDNIIISGSTDRTLKVWNAETGECIHTLYGHTSTVRCMHLHEKRVVSGSRDATLRVWDIETGQCLHVLMGHVAAVRCVQYDGRRVVSG 437
FBXW7-Canonical    CLRTLVGHTGGVWSSQMRDNIIISGSTDRTLKVWNAETGECIHTLYGHTSTVRCMHLHEKRVVSGSRDATLRVWDIETGQCLHVLMGHVAAVRCVQYDGRRVVSG 517
FBXW7-201          CLRTLVGHTGGVWSSQMRDNIIISGSTDRTLKVWNAETGECIHTLYGHTSTVRCMHLHEKRVVSGSRDATLRVWDIETGQCLHVLMGHVAAVRCVQYDGRRVVSG 517
FBXW7-204          CLRTLVGHTGGVWSSQMRDNIIISGSTDRTLKVWNAETGECIHTLYGHTSTVRCMHLHEKRVVSGSRDATLRVWDIETGQCLHVLMGHVAAVRCVQYDGRRVVSG 517
FBXW7-206          CLRTLVGHTGGVWSSQMRDNIIISGSTDRTLKVWNAETGECIHTLYGHTSTVRCMHLHEKRVVSGSRDATLRVWDIETGQCLHVLMGHVAAVRCVQYDGRRVVSG 517
IPR001810          .........................................................................................................  48
PF12937            .........................................................................................................  45
consensus          *********************************************************************************************************

logo               AYDFMVKVWDPETETCLHTLQGHTNRVYSLQFDGIHVVSGSLDTSIRVWDVETGNCIHTLTGHQSLTSGMELKDNILVSGNADSTVKIWDIKTGQCLQTLQGPNK
Pocket-Binding     .Y.........................Y.............................................................................   8
FBXW7-202          AYDFMVKVWDPETETCLHTLQGHTNRVYSLQFDGIHVVSGSLDTSIRVWDVETGNCIHTLTGHQSLTSGMELKDNILVSGNADSTVKIWDIKTGQCLQTLQGPNK 504
FBXW7-203          AYDFMVKVWDPETETCLHTLQGHTNRVYSLQFDGIHVVSGSLDTSIRVWDVETGNCIHTLTGHQSLTSGMELKDNILVSGNADSTVKIWDIKTGQCLQTLQGPNK 542
FBXW7-Canonical    AYDFMVKVWDPETETCLHTLQGHTNRVYSLQFDGIHVVSGSLDTSIRVWDVETGNCIHTLTGHQSLTSGMELKDNILVSGNADSTVKIWDIKTGQCLQTLQGPNK 622
FBXW7-201          AYDFMVKVWDPETETCLHTLQGHTNRVYSLQFDGIHVVSGSLDTSIRVWDVETGNCIHTLTGHQSLTSGMELKDNILVSGNADSTVKIWDIKTGQCLQTLQGPNK 622
FBXW7-204          AYDFMVKVWDPETETCLHTLQGHTNRVYSLQFDGIHVVSGSLDTSIRVWDVETGNCIHTLTGHQSLTSGMELKDNILVSGNADSTVKIWDIKTGQCLQTLQGPNK 622
FBXW7-206          AYDFMVKVWDPETETCLHTLQGHTNRVYSLQFDGIHVVSGSLDTSIRVWDVETGNCIHTLTGHQSLTSGMELKDNILVSGNADSTVKIWDIKTGQCLQTLQGPNK 622
IPR001810          .........................................................................................................  48
PF12937            .........................................................................................................  45
consensus          *********************************************************************************************************

logo               HQSAVTCLQFNKNFVITSSDDGTVKLWDLKTGEFIRNLVTLESGGSGGVVWRIRASNTKLVCAVGSRNGTEETKLLVLDFDVDMK
Pocket-Binding     .....................................................................................   8
FBXW7-202          HQSAVTCLQFNKNFVITSSDDGTVKLWDLKTGEFIRNLVTLESGGSGGVVWRIRASNTKLVCAVGSRNGTEETKLLVLDFDVDMK 589
FBXW7-203          HQSAVTCLQFNKNFVITSSDDGTVKLWDLKTGEFIRNLVTLESGGSGGVVWRIRASNTKLVCAVGSRNGTEETKLLVLDFDVDMK 627
FBXW7-Canonical    HQSAVTCLQFNKNFVITSSDDGTVKLWDLKTGEFIRNLVTLESGGSGGVVWRIRASNTKLVCAVGSRNGTEETKLLVLDFDVDMK 707
FBXW7-201          HQSAVTCLQFNKNFVITSSDDGTVKLWDLKTGEFIRNLVTLESGGSGGVVWRIRASNTKLVCAVGSRNGTEETKLLVLDFDVDMK 707
FBXW7-204          HQSAVTCLQFNKNFVITSSDDGTVKLWDLKTGEFIRNLVTLESGGSGGVVWRIRASNTKLVCAVGSRNGTEETKLLVLDFDVDMK 707
FBXW7-206          HQSAVTCLQFNKNFVITSSDDGTVKLWDLKTGEFIRNLVTLESGGSGGVVWRIRASNTKLVCAVGSRNGTEETKLLVLDFDVDMK 707
IPR001810          .....................................................................................  48
PF12937            .....................................................................................  45
consensus          *************************************************************************************
```

X non-conserved
X similar
X $\geq$ 50% conserved

```
@
@
@

  @
  @
logo            MDNMSITNTPTSNDACLSIVHSLMCHRQGGESETFAKRAIESLVKKLKEKKDELDSLITAITTNGAHPSKCVTIQRTLDGRLQVAGRKGFPHVIYARLWRWPDLH
IPR013019       .................SIVHSLMCHRQGGESETFAKRAIESLVKKLKEKKDELDSLITAITTNGAHPSKCVTIQRTLDGRLQVAGRKGFPHVIYARLWRWPDLH   88
IPR003619       ..............................ESETFAKRAIESLVKKLKEKKDELDSLITAITTNGAHPSKCVTIQRTLDGRLQVAGRKGFPHVIYARLWRWPDLH   75
PF03166         .........................................................................................................    0
SMAD4-205       MDNMSITNTPTSNDACLSIVHSLMCHRQGGESETFAKRAIESLVKKLKEKKDELDSLITAITTNGAHPSKCVTIQRTLDGRLQVAGRKGFPHVIYARLWRWPDLH  105
SMAD4-Canonical MDNMSITNTPTSNDACLSIVHSLMCHRQGGESETFAKRAIESLVKKLKEKKDELDSLITAITTNGAHPSKCVTIQRTLDGRLQVAGRKGFPHVIYARLWRWPDLH  105
SMAD4-201       MDNMSITNTPTSNDACLSIVHSLMCHRQGGESETFAKRAIESLVKKLKEKKDELDSLITAITTNGAHPSKCVTIQRTLDGRLQVAGRKGFPHVIYARLWRWPDLH  105
SMAD4-202       MDNMSITNTPTSNDACLSIVHSLMCHRQGGESETFAKRAIESLVKKLKEKKDELDSLITAITTNGAHPSKCVTIQRTLDGRLQVAGRKGFPHVIYARLWRWPDLH  105
IPR001132       .........................................................................................................    0
consensus       *********************************************************************************************************

logo            KNELKHVKYCQYAFDLKCDSVCVNPYHYERVVSPGIDLSGLTLQSNAPSSMMVKDEYVHDFEGQPSLSTEGHSIQTIQHPPSNRASTETYSTPALLAPSESNATS
IPR013019       KNELKHVKYCQYAFDLKCDSVCVNPYHYERVVSPGI.....................................................................  124
IPR003619       KNELKHVKYCQYAFDLKCDSVCVNP................................................................................  100
PF03166         .........................................................................................................    0
SMAD4-205       KNELKHVKYCQYAFDLKCDSVCVNPYHYERVVSPGIDLSGLTLQSNAPSSMMVKDEYVHDFEGQPSLSTEGHSIQTIQHPPSNRASTETYSTPALLAPSESNATS  210
SMAD4-Canonical KNELKHVKYCQYAFDLKCDSVCVNPYHYERVVSPGIDLSGLTLQSNAPSSMMVKDEYVHDFEGQPSLSTEGHSIQTIQHPPSNRASTETYSTPALLAPSESNATS  210
SMAD4-201       KNELKHVKYCQYAFDLKCDSVCVNPYHYERVVSPGIDLSGLTLQSNAPSSMMVKDEYVHDFEGQPSLSTEGHSIQTIQHPPSNRASTETYSTPALLAPSESNATS  210
SMAD4-202       KNELKHVKYCQYAFDLKCDSVCVNPYHYERVVSPGIDLSGLTLQSNAPSSMMVKDEYVHDFEGQPSLSTEGHSIQTIQHPPSNRASTETYSTPALLAPSESNATS  210
IPR001132       .........................................................................................................    0
consensus       *********************************************************************************************************

logo            TANFPNIPVASTSQPASILGGSHSEGLLQIASGPQPGQQQNGFTGQPATYHHNSTTTWTGSRTAPYTPNLPHHQNGHLQHHPPMPPHPGHYWPVHNELAFQPPIS
IPR013019       .........................................................................................................  124
IPR003619       .........................................................................................................  100
PF03166         .........................................................................................................    0
SMAD4-205       TANFPNIPVAST.............................................................................................  222
SMAD4-Canonical TANFPNIPVASTSQPASILGGSHSEGLLQIASGPQPGQQQNGFTGQPATYHHNSTTTWTGSRTAPYTPNLPHHQNGHLQHHPPMPPHPGHYWPVHNELAFQPPIS  315
SMAD4-201       TANFPNIPVASTSQPASILGGSHSEGLLQIASGPQPGQQQNGFTGQPATYHHNSTTTWTGSRTAPYTPNLPHHQNGHLQHHPPMPPHPGHYWPVHNELAFQPPIS  315
SMAD4-202       TANFPNIPVASTSQPASILGGSHSEGLLQIASGPQPGQQQNGFTGQPATYHHNSTTTWTGSRTAPYTPNLPHHQNGHLQHHPPMPPHPGHYWPVHNELAFQPPIS  315
IPR001132       .........................................................................................................    0
consensus       ************
```

logo 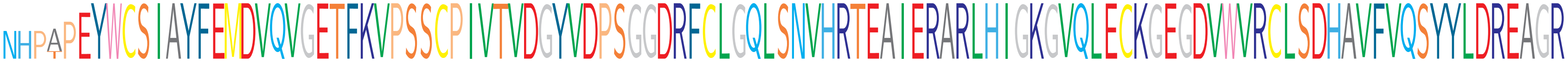

```
IPR013019         .........................................................................................................  124
IPR003619         .........................................................................................................  100
PF03166           .....EYWCSIAYFEMDVQVGETFKVPSSCPIVTVDGYVDPSGGDRFCLGQLSNVHRTEAIERARLHIGKGVQLECKGEGDVWVRCLSDHAVFVQSYYLDREAGR  100
SMAD4-205         ...TPEYWCSIAYFEMDVQVGETFKVPSSCPIVTVDGYVDPSGGDRFCLGQLSNVHRTEAIERARLHIGKGVQLECKGEGDVWVRCLSDHAVFVQSYYLDREAGR  324
SMAD4-Canonical   NHPAPEYWCSIAYFEMDVQVGETFKVPSSCPIVTVDGYVDPSGGDRFCLGQLSNVHRTEAIERARLHIGKGVQLECKGEGDVWVRCLSDHAVFVQSYYLDREAGR  420
SMAD4-201         NHPAPEYWCSIAYFEMDVQVGETFKVPSSCPIVTVDGYVDPSGGDRFCLGQLSNVHRTEAIERARLHIGKGVQLECKGEGDVWVRCLSDHAVFVQSYYLDREAGR  420
SMAD4-202         NHPAPEYWCSIAYFEMDVQVGETFKVPSSCPIVTVDGYVDPSGGDRFCLGQLSNVHRTEAIERARLHIGKGVQLECKGEGDVWVRCLSDHAVFVQSYYLDREAGR  420
IPR001132         ......YWCSIAYFEMDVQVGETFKVPSSCPIVTVDGYVDPSGGDRFCLGQLSNVHRTEAIERARLHIGKGVQLECKGEGDVWVRCLSDHAVFVQSYYLDREAGR  99
consensus             *****************************************************************************************************
```

logo 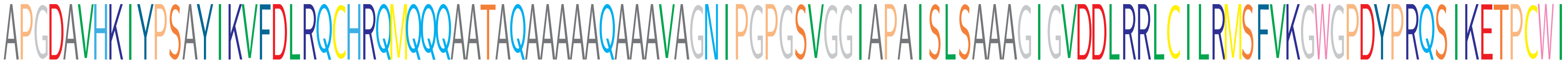

```
IPR013019         .........................................................................................................  124
IPR003619         .........................................................................................................  100
PF03166           APGDAVHKIYPSAYIKVFDLRQCHRQMQQQAATAQAAAAAQAAAVAGNIPGPGSVGGIAPAISLSAAAGIGVDDLRRLCILRMSFVKGWGPDYPRQSIKETPCWI  205
SMAD4-205         APGDAVHKIYPSAYIKVFDLRQCHRQMQQQAATAQAAAAAQAAAVAGNIPGPGSVGGIAPAISLSAAAGIGVDDLRRLCILRMSFVKGWGPDYPRQSIKETPCWI  429
SMAD4-Canonical   APGDAVHKIYPSAYIKVFDLRQCHRQMQQQAATAQAAAAAQAAAVAGNIPGPGSVGGIAPAISLSAAAGIGVDDLRRLCILRMSFVKGWGPDYPRQSIKETPCWI  525
SMAD4-201         APGDAVHKIYPSAYIKVFDLRQCHRQMQQQAATAQAAAAAQAAAVAGNIPGPGSVGGIAPAISLSAAAGIGVDDLRRLCILRMSFVKGWGPDYPRQSIKETPCWI  525
SMAD4-202         APGDAVHKIYPSAYIKVFDLRQCHRQMQQQAATAQAAAAAQAAAVAGNIPGPGSVGGIAPAISLSAAAGIGVDDLRRLCILRMSFVKGWGPDYPRQSIKETPCWI  525
IPR001132         APGDAVHKIYPSAYIKVFDLRQCHRQMQQQAATAQAAAAAQAAAVAGNIPGPGSVGGIAPAISLSAAAGIGVDDLRRLCILRMSFVKGWGPDYPRQSIKETPCWI  204
consensus         *********************************************************************************************************
```

logo 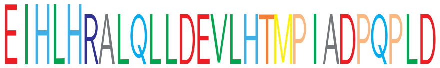

```
IPR013019         ...........................  124
IPR003619         ...........................  100
PF03166           EIHLH......................  210
SMAD4-205         EIHLHRALQLLDEVLHTMPIADPQPLD  456
SMAD4-Canonical   EIHLHRALQLLDEVLHTMPIADPQPLD  552
SMAD4-201         EIHLHRALQLLDEVLHTMPIADPQPLD  552
SMAD4-202         EIHLHRALQLLDEVLHTMPIADPQPLD  552
IPR001132         EIHLHRALQLLDEVLHTMPIADPQPLD  231
consensus         ***************************
```

X non-conserved
X similar
X ≥ 50% conserved

```
@
@
@

@
@
logo
                  MRANDALQVLGLLFSLARGSEVGNSQAVCPGTLNGLSVTGDAENQYQTLYKLYERCEVVMGNLEIVLTGHNADLSFLQWIREVTGYVLVAMNEFSTLPLPNLRVV
ERBB3-Canonical   MRANDALQVLGLLFSLARGSEVGNSQAVCPGTLNGLSVTGDAENQYQTLYKLYERCEVVMGNLEIVLTGHNADLSFLQWIREVTGYVLVAMNEFSTLPLPNLRVV  105
ERBB3-201         MRANDALQVLGLLFSLARGSEVGNSQAVCPGTLNGLSVTGDAENQYQTLYKLYERCEVVMGNLEIVLTGHNADLSFLQWIREVTGYVLVAMNEFSTLPLPNLRVV  105
ERBB3-230         ...........................................................MGNLEIVLTGHNADLSFLQWIREVTGYVLVAMNEFSTLPLPNLRVV   46
ERBB3-228         ...........................................................MGNLEIVLTGHNADLSFLQWIREVTGYVLVAMNEFSTLPLPNLRVV   46
ERBB3-227         ...........................................................MGNLEIVLTGHNADLSFLQWIREVTGYVLVAMNEFSTLPLPNLRVV   46
ERBB3-203         ...........................................................MGNLEIVLTGHNADLSFLQWIREVTGYVLVAMNEFSTLPLPNLRVV   46
IPR000494         .........................................................................................................    0
consensus                                                                    **********************************************

logo              RGTQVYDGKFAIFVMLNYNTNSSHALRQLRLTQLTEILSGGVYIEKNDKLCHMDTIDWRDIVRDRDAEIVVKDNGRSCPPCHEVCKGRCWGPGSEDCQTLTKTIC
ERBB3-213         .........................................................................................................    0
ERBB3-214         .........................................................................................................    0
IPR006211         .............................................................................CPPCHEVCKGRCWGPGSEDCQTLTKTIC   28
IPR032778         .........................................................................................................    0
ERBB3-Canonical   RGTQVYDGKFAIFVMLNYNTNSSHALRQLRLTQLTEILSGGVYIEKNDKLCHMDTIDWRDIVRDRDAEIVVKDNGRSCPPCHEVCKGRCWGPGSEDCQTLTKTIC  210
ERBB3-201         RGTQVYDGKFAIFVMLNYNTNSSHALRQLRLTQLTEILSGGVYIEKNDKLCHMDTIDWRDIVRDRDAEIVVKDNGRSCPPCHEVCKGRCWGPGSEDCQTLTKTIC  210
ERBB3-230         RGTQVYDGKFAIFVMLNYNTNSSHALRQLRLTQLTEILSGGVYIEKNDKLCHMDTIDWRDIVRDRDAEIVVKDNGRSCPPCHEVCKGRCWGPGSEDCQTLTKTIC  151
ERBB3-228         RGTQVYDGKFAIFVMLNYNTNSSHALRQLRLTQLTEILSGGVYIEKNDKLCHMDTIDWRDIVRDRDAEIVVKDNGRSCPPCHEVCKGRCWGPGSEDCQTLTKTIC  151
ERBB3-227         RGTQVYDGKFAIFVMLNYNTNSSHALRQLRLTQLTEILSGGVYIEKNDKLCHMDTIDWRDIVRDRDAEIVVKDNGRSCPPCHEVCKGRCWGPGSEDCQTLTKTIC  151
ERBB3-203         RGTQVYDGKFAIFVMLNYNTNSSHALRQLRLTQLTEILSGGVYIEKNDKLCHMDTIDWRDIVRDRDAEIVVKDNGRSCPPCHEVCKGRCWGPGSEDCQTLTKTIC  151
ERBB3-Canonical   APQCNGHCFGPNPNQCCHDECAGGCSGPQDTDCFACRHFNDSGACVPRCPQPLVYNKLTFQLEPNPHTKYQYGGVCVASCPHNFVVDQTSCVRACPPDKMEVDKN  315
ERBB3-201         APQCNGHCFGPNPNQCCHDECAGGCSGPQDTDCFACRHFNDSGACVPRCPQPLVYNKLTFQLEPNPHTKYQYGGVCVASCPHNFVVDQTSCVRACPPDKMEVDKN  315
ERBB3-230         APQCNGHCFGPNPNQCCHDECAGGCSGPQDTDCFACRHFNDSGACVPRCPQPLVYNKLTFQLEPNPHTKYQYGGVCVASCPHNFVVDQTSCVRACPPDKMEVDKN  256
ERBB3-228         APQCNGHCFGPNPNQCCHDECAGGCSGPQDTDCFACRHFNDSGACVPRCPQPLVYNKLTFQLEPNPHTKYQYGGVCVASCPHNFVVDQTSCVRACPPDKMEVDKN  256
ERBB3-227         APQCNGHCFGPNPNQCCHDECAGGCSGPQDTDCFACRHFNDSGACVPRCPQPLVYNKLTFQLEPNPHTKYQYGGVCVASCPHNFVVDQTSCVRACPPDKMEVDKN  256
ERBB3-203         APQCNGHCFGPNPNQCCHDECAGGCSGPQDTDCFACRHFNDSGACVPRCPQPLVYNKLTFQLEPNPHTKYQYGGVCVASCPHNFVVDQTSCVRACPPDKMEVDKN  256
IPR000494         .........................................................................................................    0
consensus         *********************************************************************************************************
```

```
logo             GLKMCEPCGGLCPKACEGTGSGSRFQTVDSSNIDGFVNCTKILGNLDFLITGLNGDPWHKIPALDPEKLNVFRTVREITGYLNIQSWPPHMHNFSVFSNLTTIGG
IPR032778        .........................................................................................................    0
ERBB3-Canonical  GLKMCEPCGGLCPKACEGTGSGSRFQTVDSSNIDGFVNCTKILGNLDFLITGLNGDPWHKIPALDPEKLNVFRTVREITGYLNIQSWPPHMHNFSVFSNLTTIGG  420
ERBB3-201        GLKMCEPCGGLCPKACEGTGSGSRFQTVDSSNIDGFVNCTKILGNLDFLITGLNGDPWHKIPALDPEKLNVFRTVREITGYLNIQSWPPHMHNFSVFSNLTTIGG  420
ERBB3-230        GLKMCEPCGGLCPKACEGTGSGSRFQTVDSSNIDGFVNCTKILGNLDFLITGLNGDPWHKIPALDPEKLNVFRTVREITGYLNIQSWPPHMHNFSVFSNLTTIGG  361
ERBB3-228        GLKMCEPCGGLCPKACEGTGSGSRFQTVDSSNIDGFVNCTKILGNLDFLITGLNGDPWHKIPALDPEKLNVFRTVREITGYLNIQSWPPHMHNFSVFSNLTTIGG  361
ERBB3-227        GLKMCEPCGGLCPKACEGTGSGSRFQTVDSSNIDGFVNCTKILGNLDFLITGLNGDPWHKIPALDPEKLNVFRTVREITGYLNIQSWPPHMHNFSVFSNLTTIGG  361
ERBB3-203        GLKMCEPCGGLCPKACEGTGSGSRFQTVDSSNIDGFVNCTKILGNLDFLITGLNGDPWHKIPALDPEKLNVFRTVREITGYLNIQSWPPHMHNFSVFSNLTTIGG  361
IPR000494        ..................................GFVNCTKILGNLDFLITGLNGDPWHKIPALDPEKLNVFRTVREITGYLNIQSWPPHMHNFSVFSNLTTIGG   71
consensus        *********************************************************************************************************

logo             RSLYNRGFSLLIMKNLNVTSLGFRSLKEISAGRIYISANRQLCYHHSLNWTKVLRGPTEERLDIKHNRPRRDCVAEGKVCDPLCSSGGCWGPGPGQCLSCRNYSR
ERBB3-213        .........................................................................................................    0
ERBB3-214        ................................................................................................CLSCRNYSR    9
IPR006211        .........................................................................................................  150
IPR032778        ...............................................................................CDPLCSSGGCWGPGPGQCLSCRNYSR   26
ERBB3-Canonical  RSLYNRGFSLLIMKNLNVTSLGFRSLKEISAGRIYISANRQLCYHHSLNWTKVLRGPTEERLDIKHNRPRRDCVAEGKVCDPLCSSGGCWGPGPGQCLSCRNYSR  525
ERBB3-201        RSLYNRGFSLLIMKNLNVTSLGFRSLKEISAGRIYISANRQLCYHHSLNWTKVLRGPTEERLDIKHNRPRRDCVAEGKVCDPLCSSGGCWGPGPGQCLSCRNYSR  525
ERBB3-230        RSLYNRGFSLLIMKNLNVTSLGFRSLKEISAGRIYISANRQLCYHHSLNWTKVLRGPTEERLDIKHNRPRRDCVAEGKVCDPLCSSGGCWGPGPGQCLSCRNYSR  466
ERBB3-228        RSLYNRGFSLLIMKNLNVTSLGFRSLKEISAGRIYISANRQLCYHHSLNWTKVLRGPTEERLDIKHNRPRRDCVAEGKVCDPLCSSGGCWGPGPGQCLSCRNYSR  466
ERBB3-227        RSLYNRGFSLLIMKNLNVTSLGFRSLKEISAGRIYISANRQLCYHHSLNWTKVLRGPTEERLDIKHNRPRRDCVAEGKVCDPLCSSGGCWGPGPGQCLSCRNYSR  466
ERBB3-203        RSLYNRGFSLLIMKNLNVTSLGFRSLKEISAGRIYISANRQLCYHHSLNWTKVLRGPTEERLDIKHNRPRRDCVAEGKVCDPLCSSGGCWGPGPGQCLSCRNYSR  466
IPR000494        RSLYNRGFSLLIMKNLNVTSLGFRSLKEISAGRIYISANRQLCYHHSLNW.......................................................  121
consensus        *********************************************************************************************************

logo             GGVCVTHCNFLNGEPREFAHEAECFSCHPECQPMEGTATCNGSGSDTCAQCAHFRDGPHCVSSCPHGVLGAKGPIYKYPDVQNECRPCHENCTQGCKGPELQDCL
ERBB3-213        .........................................................................................................    0
ERBB3-214        GGVCVTHCNFLNGEPREFAHEAECFSCHPECQPMEGTATCNGSGSDTCAQCAHFRDGPHCVSSCPHGVLGAKGPIYKYPDVQNECRPCHENCTQGCKGPELQDCL  114
IPR006211        .........................................................................................................  150
IPR032778        GGVCVTHCNFLNGEPREFAHEAECFSCHPECQPMEGTATCNGSGSDTCAQCAHFRDGPHCVSSCPHGVLGAKGPIYKYPDVQNECRPCHENCTQGCKGPELQDC.  130
ERBB3-Canonical  GGVCVTHCNFLNGEPREFAHEAECFSCHPECQPMEGTATCNGSGSDTCAQCAHFRDGPHCVSSCPHGVLGAKGPIYKYPDVQNECRPCHENCTQGCKGPELQDCL  630
ERBB3-201        GGVCVTHCNFLNGEPREFAHEAECFSCHPECQPMEGTATCNGSGSDTCAQCAHFRDGPHCVSSCPHGVLGAKGPIYKYPDVQNECRPCHENCTQGCKGPELQDCL  630
ERBB3-230        GGVCVTHCNFLNGEPREFAHEAECFSCHPECQPMEGTATCNGSGSDTCAQCAHFRDGPHCVSSCPHGVLGAKGPIYKYPDVQNECRPCHENCTQGCKGPELQDCL  571
ERBB3-228        GGVCVTHCNFLNGEPREFAHEAECFSCHPECQPMEGTATCNGSGSDTCAQCAHFRDGPHCVSSCPHGVLGAKGPIYKYPDVQNECRPCHENCTQGCKGPELQDCL  571
ERBB3-227        GGVCVTHCNFLNGEPREFAHEAECFSCHPECQPMEGTATCNGSGSDTCAQCAHFRDGPHCVSSCPHGVLGAKGPIYKYPDVQNECRPCHENCTQGCKGPELQDCL  571
ERBB3-203        GGVCVTHCNFLNGEPREFAHEAECFSCHPECQPMEGTATCNGSGSDTCAQCAHFRDGPHCVSSCPHGVLGAKGPIYKYPDVQNECRPCHENCTQGCKGPELQDCL  571
IPR000494        .........................................................................................................  121
consensus        *********************************************************************************************************
```

```
logo              GQTLVLIGKTHLTMALTVIAGLWIFMMLGGTFLYWRGRRIQNKRAMRRYLERGESIEPLDPSEKANKVLARIFKETELRKLKVLGSGVFGTVHKGVWIPEGESI
ERBB3-213         .........................................................................................................   0
ERBB3-214         GQTLVLIGKTHLTMALTVIAGLVVIFMMLGGTFLYWRGRRIQNKRAMRRYLERGESIEPLDPSE.........................................   178
IPR006211         .........................................................................................................   150
IPR032778         .........................................................................................................   130
ERBB3-Canonical   GQTLVLIGKTHLTMALTVIAGLVVIFMMLGGTFLYWRGRRIQNKRAMRRYLERGESIEPLDPSEKANKVLARIFKETELRKLKVLGSGVFGTVHKGVWIPEGESI   735
ERBB3-201         GQTLVLIGKTHLTMALTVIAGLVVIFMMLGGTFLYWRGRRIQNKRAMRRYLERGESIEPLDPSEKANKVLARIFKETELRKLKVLGSGVFGTVHKGVWIPEGESI   735
ERBB3-230         GQTLVLIGKTHLTMALTVIAGLVVIFMMLGGTFLYWRGRRIQNKRAMRRYLERGESIEPLDPSEKANKVLARIFKETELRKLKVLGSGVFGTVHKGVWIPEGESI   676
ERBB3-228         GQTLVLIGKTHLTMALTVIAGLVVIFMMLGGTFLYWRGRRIQNKRAMRRYLERGESIEPLDPSEKANKVLARIFKETELRKLKVLGSGVFGTVHKGVWIPEGESI   676
ERBB3-227         GQTLVLIGKTHLTMALTVIAGLVVIFMMLGGTFLYWRGRRIQNKRAMRRYLERGESIEPLDPSEKANKVLARIFKETELRKLKVLGSGVFGTVHKGVWIPEGESI   676
ERBB3-203         GQTLVLIGKTHLTMALTVIAGLVVIFMMLGGTFLYWRGRRIQNKRAMRRYLERGESIEPLDPSEKANKVLARIFKETELRKLKVLGSGVFGTVHKGVWIPEGESI   676
IPR000494         .........................................................................................................   121
consensus         *********************************************************************************************************

logo              KIPVCIKVIEDKSGRQSFQAVTDHMLAIGSLDHAHIVRLLGLCPGSSLQLVTQYLPLGSLLDHVRQHRGALGPQLLLNWGVQIAKGMYYLEEHGMVHRNLAARNV
ERBB3-213         .........................................................................................................   0
ERBB3-214         .........................................................................................................   178
IPR006211         .........................................................................................................   150
IPR032778         .........................................................................................................   130
ERBB3-Canonical   KIPVCIKVIEDKSGRQSFQAVTDHMLAIGSLDHAHIVRLLGLCPGSSLQLVTQYLPLGSLLDHVRQHRGALGPQLLLNWGVQIAKGMYYLEEHGMVHRNLAARNV   840
ERBB3-201         KIPVCIKVIEDKSGRQSFQAVTDHMLAIGSLDHAHIVRLLGLCPGSSLQLVTQYLPLGSLLDHVRQHRGALGPQLLLNWGVQIAKGMYYLEEHGMVHRNLAARNV   840
ERBB3-230         KIPVCIKVIEDKSGRQSFQAVTDHMLAIGSLDHAHIVRLLGLCPGSSLQLVTQYLPLGSLLDHVRQHRGALGPQLLLNWGVQIAKGMYYLEEHGMVHRNLAARNV   781
ERBB3-228         KIPVCIKVIEDKSGRQSFQAVTDHMLAIGSLDHAHIVRLLGLCPGSSLQLVTQYLPLGSLLDHVRQHRGALGPQLLLNWGVQIAKGMYYLEEHGMVHRNLAARNV   781
ERBB3-227         KIPVCIKVIEDKSGRQSFQAVTDHMLAIGSLDHAHIVRLLGLCPGSSLQLVTQYLPLGSLLDHVRQHRGALGPQLLLNWGVQIAKGMYYLEEHGMVHRNLAARNV   781
ERBB3-203         KIPVCIKVIEDKSGRQSFQAVTDHMLAIGSLDHAHIVRLLGLCPGSSLQLVTQYLPLGSLLDHVRQHRGALGPQLLLNWGVQIAKGMYYLEEHGMVHRNLAARNV   781
IPR000494         .........................................................................................................   121
consensus         *********************************************************************************************************

logo              LLKSPSQVQVADFGVADLLPPDDKQLLYSEAKTPIKWMALESIHFGKYTHQSDVWSYGVTVWELMTFGAEPYAGLRLAEVPDLLEKGERLAQPQICTIDVYMVMV
ERBB3-213         .....................................MPMSTIL...PVTSPCLPATGVTVWELMTFGAEPYAGLRLAEVPDLLEKGERLAQPQICTIDVYMVMV   65
ERBB3-214         ...............................KTPIKWMALESIHFGKYTHQSDVWSYGVTVWELMTFGAEPYAGLRLAEVPDLLEKGERLAQPQICTIDVYMVMV   252
IPR006211         .........................................................................................................   150
IPR032778         .........................................................................................................   130
ERBB3-Canonical   LLKSPSQVQVADFGVADLLPPDDKQLLYSEAKTPIKWMALESIHFGKYTHQSDVWSYGVTVWELMTFGAEPYAGLRLAEVPDLLEKGERLAQPQICTIDVYMVMV   945
ERBB3-201         LLKSPSQVQVADFGVADLLPPDDKQLLYSEAKTPIKWMALESIHFGKYTHQSDVWSYGVTVWELMTFGAEPYAGLRLAEVPDLLEKGERLAQPQICTIDVYMVMV   945
ERBB3-230         LLKSPSQVQVADFGVADLLPPDDKQLLYSEAKTPIKWMALESIHFGKYTHQSDVWSYGVTVWELMTFGAEPYAGLRLAEVPDLLEKGERLAQPQICTIDVYMVMV   886
ERBB3-228         LLKSPSQVQVADFGVADLLPPDDKQLLYSEAKTPIKWMALESIHFGKYTHQSDVWSYGVTVWELMTFGAEPYAGLRLAEVPDLLEKGERLAQPQICTIDVYMVMV   886
ERBB3-227         LLKSPSQVQVADFGVADLLPPDDKQLLYSEAKTPIKWMALESIHFGKYTHQSDVWSYGVTVWELMTFGAEPYAGLRLAEVPDLLEKGERLAQPQICTIDVYMVMV   886
ERBB3-203         LLKSPSQVQVADFGVADLLPPDDKQLLYSEAKTPIKWMALESIHFGKYTHQSDVWSYGVTVWELMTFGAEPYAGLRLAEVPDLLEKGERLAQPQICTIDVYMVMV   886
IPR000494         .........................................................................................................   121
consensus         *********************************************************************************************************
```

```
logo             KCWMIDENIRPTFKELANEFTRMARDPPRYLVIKRESGPGIAPGPEPHGLTNKKLEEVELEPELDLDLDLEAEEDNLATTTLGSALSLPVGTLNRPRGSQSLLSP
ERBB3-213        KCWMIDENIRPTFKELANEFTRMARDPPRYLVIKRESGPGIAPGPEPHGLTNKKLEEVELEPELDLDLDLEAEEDNLATTTLGSALSLPVGTLNRPRGSQSLLSP  170
ERBB3-214        KCWMIDENIRPTFKELANEFTRMARDPPRYLVIKRESGPGIAPGPEPHGLTNKKLEEVELEPELDLDLDLEAEEDNLATTTLGSALSLPVGTLNRPRGSQSLLSP  357
IPR006211        .........................................................................................................  150
IPR032778        .........................................................................................................  130
ERBB3-Canonical  KCWMIDENIRPTFKELANEFTRMARDPPRYLVIKRESGPGIAPGPEPHGLTNKKLEEVELEPELDLDLDLEAEEDNLATTTLGSALSLPVGTLNRPRGSQSLLSP  1050
ERBB3-201        KCWMIDENIRPTFKELANEFTRMARDPPRYLVIKRESGPGIAPGPEPHGLTNKKLEEVELEPELDLDLDLEAEEDNLATTTLGSALSLPVGTLNRPRGSQSLLSP  1050
ERBB3-230        KCWMIDENIRPTFKELANEFTRMARDPPRYLVIKRESGPGIAPGPEPHGLTNKKLEEVELEPELDLDLDLEAEEDNLATTTLGSALSLPVGTLNRPRGSQSLLSP  991
ERBB3-228        KCWMIDENIRPTFKELANEFTRMARDPPRYLVIKRESGPGIAPGPEPHGLTNKKLEEVELEPELDLDLDLEAEEDNLATTTLGSALSLPVGTLNRPRGSQSLLSP  991
ERBB3-227        KCWMIDENIRPTFKELANEFTRMARDPPRYLVIKRESGPGIAPGPEPHGLTNKKLEEVELEPELDLDLDLEAEEDNLATTTLGSALSLPVGTLNRPRGSQSLLSP  991
ERBB3-203        KCWMIDENIRPTFKELANEFTRMARDPPRYLVIKRESGPGIAPGPEPHGLTNKKLEEVELEPELDLDLDLEAEEDNLATTTLGSALSLPVGTLNRPRGSQSLLSP  991
IPR000494        .........................................................................................................  121
consensus        *********************************************************************************************************

logo             SSGYMPMNQGNLGESCQESAVSGSSERCPRPVSLHPMPRGCLASESSEGHVTGSEAELQEKVSMCRSRSRSRSPRPRGDSAYHSQRHSLLTPVTPLSPPGLEEED
ERBB3-213        SSGYMPMNQGNLGESCQESAVSGSSERCPRPVSLHPMPRGCLASESSEGHVTGSEAELQEKVSMCRSRSRSRSPRPRGDSAYHSQRHSLLTPVTPLSPPGLEEED  275
ERBB3-214        SSGYMPMNQGNLGESCQESAVSGSSERCPRPVSLHPMPRGCLASESSEGHVTGSEAELQEKVSMCRSRSRSRSPRPRGDSAYHSQRHSLLTPVTPLSPPGLEEED  462
IPR006211        .........................................................................................................  150
IPR032778        .........................................................................................................  130
ERBB3-Canonical  SSGYMPMNQGNLGESCQESAVSGSSERCPRPVSLHPMPRGCLASESSEGHVTGSEAELQEKVSMCRSRSRSRSPRPRGDSAYHSQRHSLLTPVTPLSPPGLEEED  1155
ERBB3-201        SSGYMPMNQGNLGESCQESAVSGSSERCPRPVSLHPMPRGCLASESSEGHVTGSEAELQEKVSMCRSRSRSRSPRPRGDSAYHSQRHSLLTPVTPLSPPGLEEED  1155
ERBB3-230        SSGYMPMNQGNLGESCQESAVSGSSERCPRPVSLHPMPRGCLASESSEGHVTGSEAELQEKVSMCRSRSRSRSPRPRGDSAYHSQRHSLLTPVTPLSPPGLEEED  1096
ERBB3-228        SSGYMPMNQGNLGESCQESAVSGSSERCPRPVSLHPMPRGCLASESSEGHVTGSEAELQEKVSMCRSRSRSRSPRPRGDSAYHSQRHSLLTPVTPLSPPGLEEED  1096
ERBB3-227        SSGYMPMNQGNLGESCQESAVSGSSERCPRPVSLHPMPRGCLASESSEGHVTGSEAELQEKVSMCRSRSRSRSPRPRGDSAYHSQRHSLLTPVTPLSPPGLEEED  1096
ERBB3-203        SSGYMPMNQGNLGESCQESAVSGSSERCPRPVSLHPMPRGCLASESSEGHVTGSEAELQEKVSMCRSRSRSRSPRPRGDSAYHSQRHSLLTPVTPLSPPGLEEED  1096
IPR000494        .........................................................................................................  121
consensus        *********************************************************************************************************

logo             VNGYVMPDTHLKGTPSSREGTLSSVGLSSVLGTEEEDEDEEYEYMNRRRRHSPPHPPRPSSLEELGYEYMDVGSDLSASLGSTQSCPLHPVPIMPTAGTTPDEDY
ERBB3-213        VNGYVMPDTHLKGTPSSREGTLSSVGLSSVLGTEEEDEDEEYEYMNRRRRHSPPHPPRPSSLEELGYEYMDVGSDLSASLGSTQSCPLHPVPIMPTAGTTPDEDY  380
ERBB3-214        VNGYVMPDTHLKGTPSSREGTLSSVGLSSVLGTEEEDEDEEYEYMNRRRRHSPPHPPRPSSLEELGYEYMDVGSDLSASLGSTQSCPLHPVPIMPTAGTTPDEDY  567
IPR006211        .........................................................................................................  150
IPR032778        .........................................................................................................  130
ERBB3-Canonical  VNGYVMPDTHLKGTPSSREGTLSSVGLSSVLGTEEEDEDEEYEYMNRRRRHSPPHPPRPSSLEELGYEYMDVGSDLSASLGSTQSCPLHPVPIMPTAGTTPDEDY  1260
ERBB3-201        VNGYVMPDTHLKGTPSSREGTLSSVGLSSVLGTEEEDEDEEYEYMNRRRRHSPPHPPRPSSLEELGYEYMDVGSDLSASLGSTQSCPLHPVPIMPTAGTTPDEDY  1260
ERBB3-230        VNGYVMPDTHLKGTPSSREGTLSSVGLSSVLGTEEEDEDEEYEYMNRRRRHSPPHPPRPSSLEELGYEYMDVGSDLSASLGSTQSCPLHPVPIMPTAGTTPDEDY  1201
ERBB3-228        VNGYVMPDTHLKGTPSSREGTLSSVGLSSVLGTEEEDEDEEYEYMNRRRRHSPPHPPRPSSLEELGYEYMDVGSDLSASLGSTQSCPLHPVPIMPTAGTTPDEDY  1201
ERBB3-227        VNGYVMPDTHLKGTPSSREGTLSSVGLSSVLGTEEEDEDEEYEYMNRRRRHSPPHPPRPSSLEELGYEYMDVGSDLSASLGSTQSCPLHPVPIMPTAGTTPDEDY  1201
ERBB3-203        VNGYVMPDTHLKGTPSSREGTLSSVGLSSVLGTEEEDEDEEYEYMNRRRRHSPPHPPRPSSLEELGYEYMDVGSDLSASLGSTQSCPLHPVPIMPTAGTTPDEDY  1201
IPR000494        .........................................................................................................  121
consensus        *********************************************************************************************************
```

```
logo              EYMNRQRDGGGPGGDYAAMGACPASEQGYEEMRAFQGPGHQAPHVHYARLKTLRSLEATDSAFDNPDYWHSRLFPKANAQRT
ERBB3-213         EYMNRQRDGGGPGGDYAAMGACPASEQGYEEMRAFQGPGHQAPHVHYARLKTLRSLEATDSAFDNPDYWHSRLFPKANAQRT  462
ERBB3-214         EYMNRQRDGGGPGGDYAAMGACPASEQGYEEMRAFQGPGHQAPHVHYARLKTLRSLEATDSAFDNPDYWHSRLFPKANAQRT  649
IPR006211         ..................................................................................  150
IPR032778         ..................................................................................  130
ERBB3-Canonical   EYMNRQRDGGGPGGDYAAMGACPASEQGYEEMRAFQGPGHQAPHVHYARLKTLRSLEATDSAFDNPDYWHSRLFPKANAQRT 1342
ERBB3-201         EYMNRQRDGGGPGGDYAAMGACPASEQGYEEMRAFQGPGHQAPHVHYARLKTLRSLEATDSAFDNPDYWHSRLFPKANAQRT 1342
ERBB3-230         EYMNRQRDGGGPGGDYAAMGACPASEQGYEEMRAFQGPGHQAPHVHYARLKTLRSLEATDSAFDNPDYWHSRLFPKANAQRT 1283
ERBB3-228         EYMNRQRDGGGPGGDYAAMGACPASEQGYEEMRAFQGPGHQAPHVHYARLKTLRSLEATDSAFDNPDYWHSRLFPKANAQRT 1283
ERBB3-227         EYMNRQRDGGGPGGDYAAMGACPASEQGYEEMRAFQGPGHQAPHVHYARLKTLRSLEATDSAFDNPDYWHSRLFPKANAQRT 1283
ERBB3-203         EYMNRQRDGGGPGGDYAAMGACPASEQGYEEMRAFQGPGHQAPHVHYARLKTLRSLEATDSAFDNPDYWHSRLFPKANAQRT 1283
IPR000494         ..................................................................................  121
consensus         **********************************************************************************
```

X non-conserved
X similar
X ≥ 50% conserved
