## Supplementary File 4 for "In Silico Analysis of Drug Off-Target Effects on Diverse Isoforms of Cervical Cancer for Enhanced Therapeutic Strategies"

### Molecular Docking Results

#### 1. KRAS Protein

| Drugs | KRAS-Canonical | KRAS-201 | KRAS-202 | KRAS-205 | KRAS-214 |
|---|---|---|---|---|---|
| AZD-4785 | -6.9 | -7.3 | -7.2 | -7 | -7.2 |
| AZD-8835 | -8.3 | -8.2 | -8.2 | -8.7 | -7.9 |
| CC-223 | -7.2 | -7.7 | -7.5 | -7.5 | -7.7 |
| PD-0325901 | -7.7 | -7.3 | -6.9 | -6.9 | -7.4 |
| RIDAFOROLIMUS | -10.1 | -10.2 | -9.7 | -9.6 | -10.6 |
| SELUMETINIB | -7.2 | -7.2 | -7.6 | -7.7 | -7.1 |
| TRAMETINIB | -8.2 | -9 | -7.9 | -9.6 | -9 |

| Drugs | KRAS-213 | KRAS-210 | KRAS-207 | KRAS-203 | KRAS-204 |
|---|---|---|---|---|---|
| AZD-4785 | -6.4 | -7.9 | -5.8 | -5.7 | -4.7 |
| AZD-8835 | -7.7 | -8.5 | -6.5 | -6.3 | -4.9 |
| CC-223 | -7.3 | -7.7 | -6.1 | -6.1 | -4.5 |
| PD-0325901 | -6.3 | -6.8 | -6.1 | -5.4 | -4.3 |
| RIDAFOROLIMUS | -10.7 | -9.6 | -9.3 | -8.8 | -5.3 |
| SELUMETINIB | -6.6 | -7.4 | -5.8 | -5.9 | -4.8 |
| TRAMETINIB | -8.6 | -8.9 | -7 | -6.9 | -5.5 |

#### 2. PIK3CA Protein

| FDA Drugs | PIK3CA-Canonical | PIK3CA-201 | PIK3CA-205 | PIK3CA-203 | PIK3CA-204 |
|---|---|---|---|---|---|
| ALPELISIB | -8.9 | -8.8 | -9.5 | -7.5 | -7.7 |
| BUPARLISIB | -8.6 | -8.2 | -8.1 | -6.2 | -6.4 |
| CAPIVASERTIB | -9.5 | -9.6 | -8.9 | -6.6 | -6.6 |
| INK-1117 | -9 | -9 | -9 | -6.8 | -6.8 |
| SERABELISIB | -8.9 | -9.1 | -9 | -6.8 | -6.8 |
| TASELISIB | -9.5 | -9.7 | -8.2 | -7.2 | -6.9 |
| Temsirolimus | -11.5 | -11.6 | -10.2 | -9 | -9 |
| Trastuzumab | -10.5 | -9.6 | -9.6 | -6.8 | -7.7 |
| CC-223 | -8.6 | -8.3 | -8 | -6.4 | -6.4 |

#### 3. ERBB3 Protein

| Drugs | ERBB3_Canonical | ERBB3-222 | ERBB3-215 | ERBB3-207 | ERBB3-213 |
|---|---|---|---|---|---|
| Pertuzumab | -5.4 | -4.4 | -5.7 | -5.7 | 6.3 |
| SAPITINIB | -6.8 | -5.5 | -7 | -7.3 | -7.3 |
| Trastuzumab | -9.4 | -7 | -8 | -6.2 | -7.2 |

| Drugs | ERBB3-202 | ERBB3-209 |
|---|---|---|
| Pertuzumab | -4.7 | -5.4 |
| SAPITINIB | -6.2 | -6.6 |
| Trastuzumab | -7.6 | -7.9 |

#### 4. SMAD4 Protein

| Drugs | SMAD4-Canonical | SMAD4-201 | SMAD4-202 | SMAD4-205 | SMAD4-206 |
|---|---|---|---|---|---|
| ALECTINIB | -7.7 | -8.6 | -9.4 | -9.9 | -8.3 |
| CRIZOTINIB | -7.1 | -7.2 | -9.8 | -8.7 | -8 |
| FLUOROURACIL | -4.7 | -4.7 | -4.9 | -4.8 | -5 |
| GEMCITABINE | -6.1 | -5.3 | -7 | -5.4 | -7.2 |
| IRINOTECAN | -9.2 | -9.3 | -10.1 | -9.5 | -9.5 |
| Lysine | -4.9 | -4.7 | -4.6 | -4.2 | -4.7 |
| SAPANISERTIB | -7 | | -8.6 | -7.1 | -7.2 |

| Drugs | SMAD4-207 | SMAD4-210 |
|---|---|---|
| ALECTINIB | -8.6 | -7.5 |
| CRIZOTINIB | -8 | -6.8 |
| FLUOROURACIL | -4.3 | -4.4 |
| GEMCITABINE | -5.6 | -5.1 |
| IRINOTECAN | -8.9 | -8.4 |
| Lysine | -4.3 | -4.1 |
| SAPANISERTIB | -7 | -6.4 |

#### 5. FBXW7 Protein

| Drugs | FBXW7-Canonical | FBXW7-202 | FBXW7-212 | FBXW7-213 | FBXW7-214 |
|---|---|---|---|---|---|
| AR-42 | -7.2 | -7.1 | -5.7 | -5.7 | -5.3 |
| BELINOSTAT | -7.4 | -7.2 | -5.3 | -5.3 | -5.9 |
| ENTINOSTAT | -7.9 | -7.8 | -5.7 | -5.7 | -6 |
| REGORAFENIB | -8.3 | -8.3 | -5.9 | -6.3 | -6.5 |
| Sirolimus | -10.5 | -10.4 | -7.4 | -7 | -6.9 |
| VORINOSTAT | -6.3 | -6.1 | -4.7 | -4.7 | -5 |

| Drugs | FBXW7-215 | FBXW7-203 |
|---|---|---|
| AR-42 | -5.7 | -6.9 |
| BELINOSTAT | -5.7 | -7.3 |
| ENTINOSTAT | -5.9 | -7.5 |
| REGORAFENIB | -6.3 | -9 |
| Sirolimus | -6.7 | -9.2 |
| VORINOSTAT | -4.8 | -6.4 |
